# Optical metabolic imaging of the tricarboxylic acid cycle

**DOI:** 10.64898/2026.01.28.702395

**Authors:** Rahuljeet S. Chadha, Philip A. Kocheril, Adrian Colazo, Edmund D. Kapelczak, Berea Suen, Ziguang Yang, Tara A. TeSlaa, Lu Wei

**Affiliations:** Division of Chemistry and Chemical Engineering, California Institute of Technology, Pasadena, CA 91125, USA; Department of Molecular and Medical Pharmacology, David Geffen School of Medicine at UCLA, Los Angeles, CA 90095, USA

**Author notes:** (L.W.). These authors contributed equally (Rahuljeet S. Chadha, Philip A. Kocheril, Adrian Colazo).

## Abstract

The tricarboxylic acid (TCA) cycle lies at the core of cellular metabolism, integrating energy production, biosynthesis and redox homeostasis, yet direct quantitative imaging of its activity in living systems remains challenging. Here we introduce MATRIX-SRS (Metabolic Activity TRacing of the trIcarboXylic acid cycle by Stimulated Raman Scattering microscopy), a platform enabling spatially resolved quantification of TCA-linked metabolism in live cells. Using emerging deuterium-labeled probes, MATRIX-SRS visualizes subcellular TCA-associated carbon-deuterium bonds in live cancer cells and neurons. We then integrate density functional theory, reaction network mapping, and hyperspectral MATRIX-SRS to construct a robust *in situ* metabolic quantification pipeline. Integrating MATRIX-SRS with isotope-tracing mass spectrometry, we reveal a global attenuation of TCA activity during epithelial-to-mesenchymal transition, providing deep molecular insights. Applying this framework, we further quantify changes in deuterium-labeled biomass in absolute concentrations for the first time, under native and drug-treated conditions, establishing a generalizable foundation for live quantitative spatial metabolomics.

## Introduction

The TCA cycle, also known as the Krebs cycle or the citric acid cycle, is regarded as the central metabolic engine of cells.^1^ Beyond their role in energy production, TCA intermediates function as precursors for biomacromolecule synthesis and are replenished through anaplerotic reactions to sustain the metabolic balance in cells.^2^ However, dysregulation of TCA cycle intermediates, often resulting from mutations in their regulatory enzymes, has been implicated in neurodegenerative diseases such as Alzheimer’s disease^3,4^ and Parkinson’s disease,^5^ metabolic disorders such as metabolic-associated fatty liver disease,^6^ and various cancers.^7,8^

Significant advancements have been made toward visualizing metabolites in biological systems, but the enabling methods each have intrinsic limitations. Fluorescence microscopy allows ultrasensitive dynamic visualization of metabolites at the subcellular level but may perturb their native behavior, since it usually requires labeling with bulky fluorophores that are much larger than the metabolites themselves. Genetically encodable biosensors are highly compatible with live cells, but they require specific expression that may perturb native cellular states, can exhibit bias from other environmental factors (such as pH, temperature, and viscosity),^9^ and are limited to only a handful of targets.^10^ Mass spectrometry imaging allows spatial mapping with high chemical specificity but remains destructive and hence incompatible for live-cell analysis.^11^ Positron emission tomography and magnetic resonance imaging (MRI) are highly biocompatible and non-invasive but have not yet reached subcellular resolution and still suffer from poor sensitivity and metabolic resolvability.^12,13^ Therefore, there is a critical need for tools that would enable quantitative subcellular mapping of TCA-linked metabolic activity.

Stimulated Raman scattering (SRS) microscopy has emerged as a powerful live-cell metabolic imaging platform over the past decade. Leveraging the ∼10^8^-fold stimulated emission signal amplification compared to spontaneous Raman scattering, SRS has achieved imaging sensitivity of small metabolites down to the µM level, with a spatial resolution of ∼400 nm and imaging speeds up to video rate. SRS has allowed label-free imaging by targeting the unique vibrations of chemical bonds from endogenous metabolites.^14–18^ With small vibrationally labeled (e.g., deuterium-labeled) metabolites, metabolic dynamics can be further probed by hyperspectral SRS (hSRS) while maintaining high biomolecular fidelity, owing to the minimal change of physico-chemical properties of the molecules from small vibrational tags. Notably, by coupling SRS with deuterated glucose (i.e., d_7_-glucose) labeling, glucose metabolism has been investigated for assessing newly synthesized biomass, including lipids, proteins, nucleic acids (DNA and RNA), and glycogen (each with unique Raman spectra), from live cells to animals.^19– 22^ While SRS studies with glucose have revealed new insights into cellular metabolism, studying other key metabolites involved in the TCA cycle with SRS remains challenging, mostly due to their low intracellular abundance, transient nature, and chemical heterogeneity.^23^ Another limitation of current spectral interpretations of metabolized biomolecular species is their largely empirical nature and limited resolvability. A physically grounded Raman spectral analysis would therefore substantially improve both precision and quantitative accuracy over the prior state-of-the-art.^19–22^

In this work, we describe MATRIX-SRS, Metabolic Activity TRacing of the trIcarboXylic acid cycle via SRS microscopy (**Fig. 1**). Through careful optimization, we first expand the palette of TCA probes accessible to metabolic SRS imaging using deuterium labeling, enabling visualization of TCA-associated metabolic activity from glucose, glutamine, lactate, acetate, pyruvate, and succinate in live neurons and cancer cells (**Fig. 1a**). With spectroscopic analysis, we demonstrate that our TCA probes are compatible with multiplex imaging applications for simultaneous metabolic profiling. We next demonstrate that these probes reveal globally altered metabolism underlying epithelial-to-mesenchymal transition (EMT), a process whose metabolic basis remains highly elusive. We further dissect carbon flow into the TCA cycle by focusing on pyruvate, which is uniquely positioned at a branching point between the TCA cycle, glycolysis/gluconeogenesis, amino acid synthesis, and *de novo* lipogenesis (DNL). To gain deeper molecular-level insights, we complement our MATRIX-SRS imaging with stable isotope-labeled (SIL) tracing using liquid chromatography-mass spectrometry (LC-MS), observing excellent quantitative agreement in metabolic trends in EMT. Based on these results, we develop a theoretical foundation for quantitative interpretation of biomass contributions derived from deuterated metabolites, using physics-based modeling built on density functional theory (DFT) (**Fig. 1b**). This model provides a general and accurate framework for *in situ* compositional profiling, significantly transcending the conventional empirical assignments from SRS. Finally, through pharmacological perturbation, we assess the biomass concentrations derived from d_3_-pyruvate incorporation, linking live-cell MATRIX-SRS imaging with biochemical flux measurements. Integrating experiments, theory, and modeling, MATRIX-SRS provides a unique live-cell optical strategy toward intracellular quantitative spatial metabolomics.

**Figure 1.**
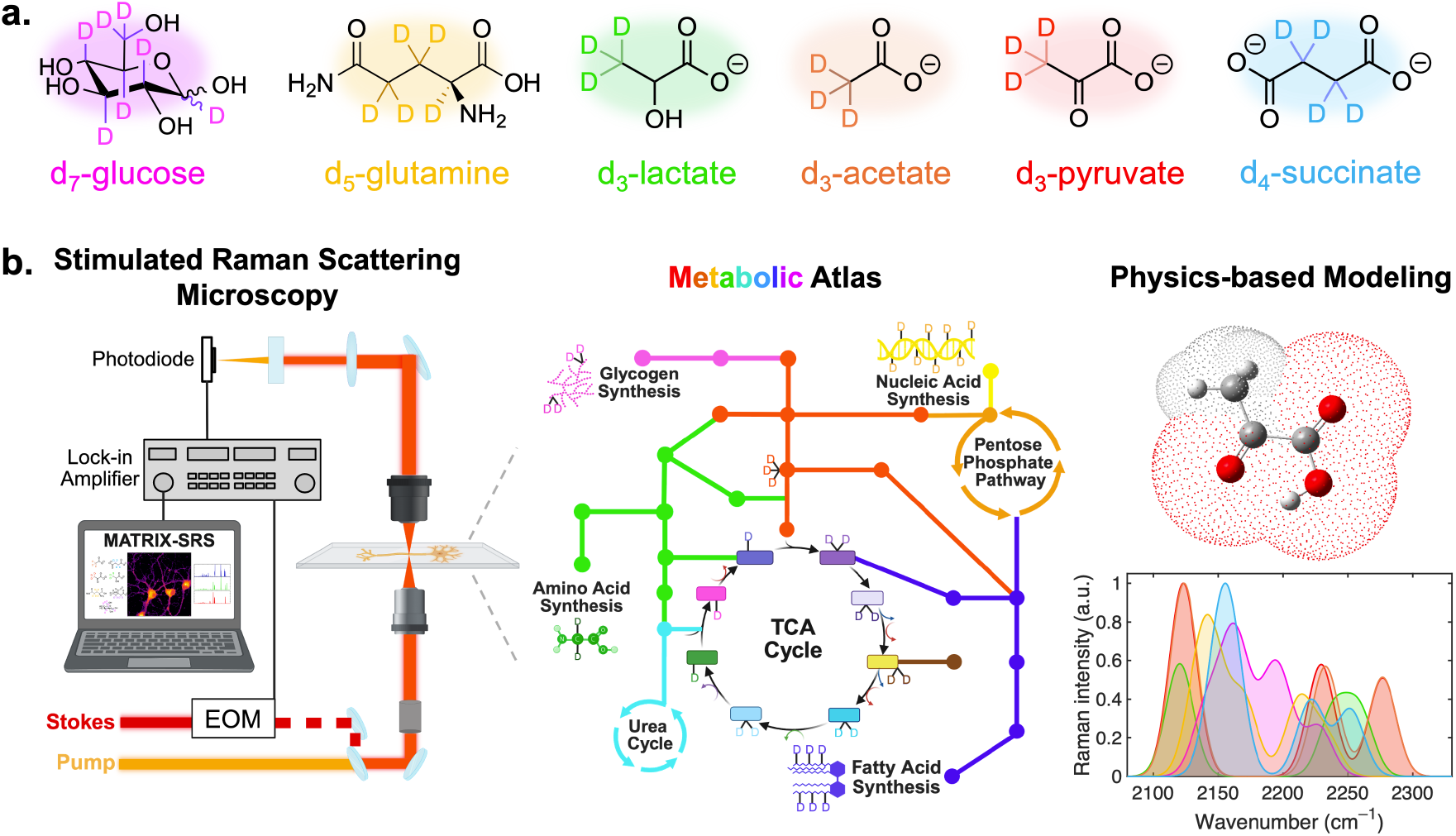
MATRIX-SRS workflow. **(a)** Chemical structures of TCA-linked Raman probes; **(b)** Left to right: schematic of the stimulated Raman scattering (SRS) microscopy setup; metabolic atlas tracing using deuterium-labeled probes; and physics-based modeling with density functional theory (DFT) for Raman peak assignments and quantification.

## Results

### I. Visualization of TCA cycle metabolism in live cells using MATRIX-SRS

We first screened stable and non-toxic TCA-related deuterium probes compatible with cell culture, from established d_7_-glucose (d_7_-glc)^19–22^ to emerging TCA probes including d_5_-glutamine (d_5_-gln),^22,24^ d_3_-lactate,^22^ d_3_-acetate, d_3_-pyruvate (d_3_-pyr), and d_4_-succinate (**Fig. 1a** and **Fig. S1**). Recently, SRS imaging of lactate and glutamine was demonstrated in gluconeogenic conditions,^22^ but our current work demonstrates quantitative imaging of these probes under native conditions, fueling new insights. After incubation for 72 hours, we directly visualized the biomass produced from each metabolite at the subcellular level with SRS in live HeLa cells (**Fig. 2a**) and live primary neuronal co-cultures (**Fig. 2b**), demonstrating the broad applicability of MATRIX-SRS imaging. We also successfully imaged d_4_-citrate in live HeLa cells (**Fig. S2**) but did not achieve similar image quality in neurons, highlighting the need for careful probe selection and optimization to obtain robust imaging results (**Fig. 2a-b**).

**Figure 2.**
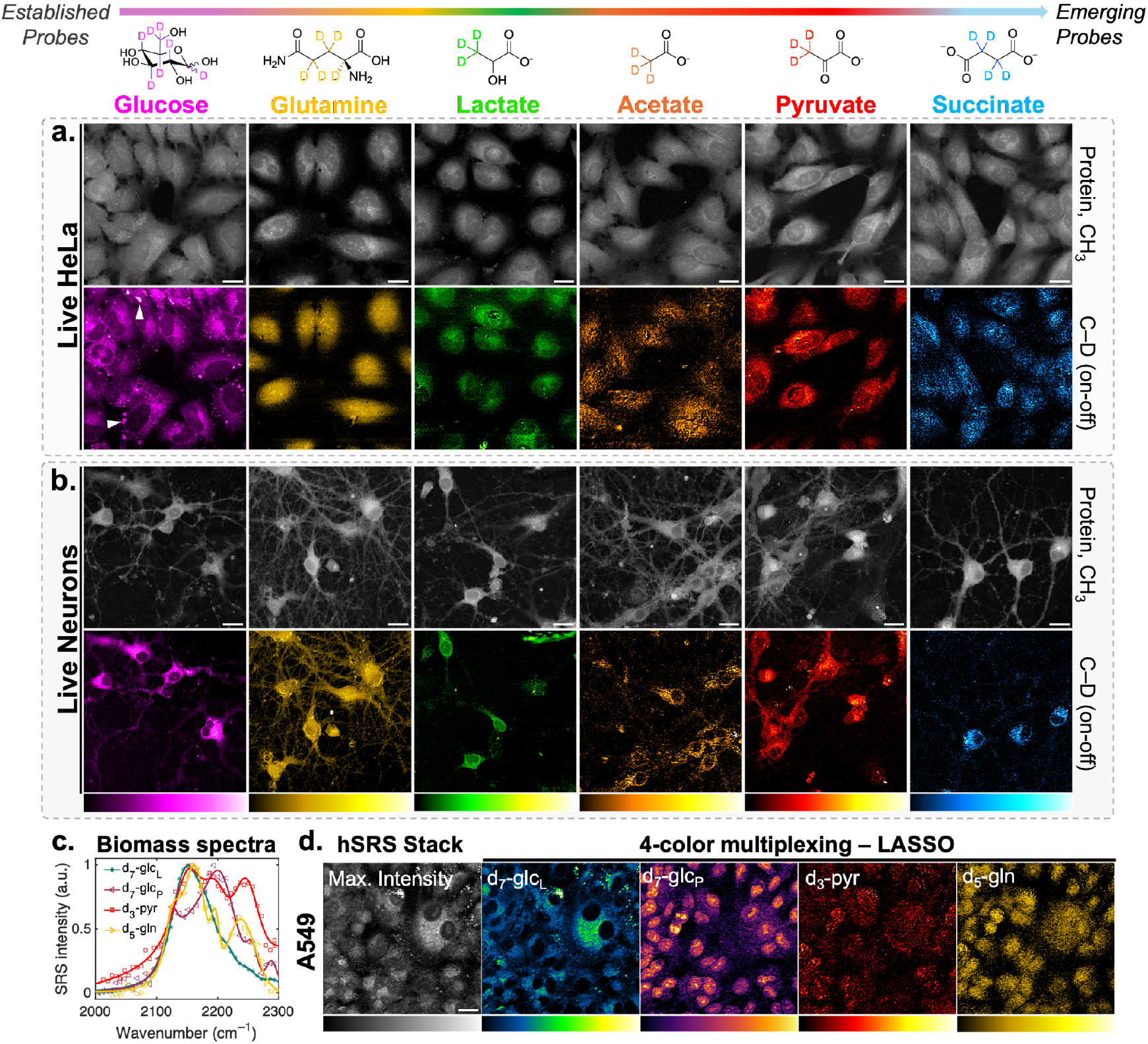
Live-cell subcellular metabolic imaging of TCA-linked activity using MATRIX-SRS. SRS images targeted at the –CH_3_ channel (protein, 2940 cm^−1^) and C–D (on-off) for six metabolic probes: d_7_-glucose, d_5_-glutamine, d_3_-lactate. d_3_-acetate, d_3_-pyruvate and d_4_-succinate in live **(a)** HeLa and **(b)** neuronal co-cultures. White arrowheads indicate subcellular glycogen pools. **(c)** Normalized hSRS spectra of the biomass from i) d_7_-glucose-derived lipids (d_7_-glc_L_; blue), ii) d_7_-glucose-derived proteins (d_7_-glc_P_; purple), iii) d_3_-pyruvate (d_3_-pyr; red), and iv) d_5_-glutamine (d_5_-gln; yellow)**; (d)** Four-color multiplex imaging of glucose, glutamine and pyruvate metabolism in A549 cells using MATRIX-SRS and LASSO unmixing; Scale bar: 20 µm.

Interestingly, SRS images also revealed distinct spatial patterns between these cell types. For instance, consistent with prior reports,^21^ we found subcellular glycogen reservoirs in live HeLa cells, but not in neurons. Additionally, in HeLa, d_3_-acetate and d_4_-succinate imaging showed a largely homogenous distribution across the entire cell, whereas in the neuronal co-culture, C–D signal from these probes was predominantly cytoplasmic. In contrast, d_3_-lactate imaging showed appreciable biomass in neuronal nuclei, demonstrating that, despite being closely chemically related, each probe can report unique subcellular metabolic information.

Since different probes can report on different metabolic processes and dynamics, complex metabolic interactions are best visualized by multiplex imaging of multiple species. It is well established that molecular vibrations act as unique ‘fingerprints,’ and using the frequency-dependence of SRS signal (i.e., hSRS) allows for direct, *in situ* profiling of these spectral fingerprints. After measuring hSRS spectra of A549 cells treated individually with each of our TCA probes, we indeed observe unique spectral fingerprints in the biomass derived from each probe (**Fig. 2c**). These biomass spectra are also substantially different from the Raman spectra of the probes in solution (**Fig. S1**), as evidence that the probes have been significantly metabolized and converted into downstream biomolecular species.

Although the biomass spectra are significantly overlapped (**Fig. 2c**), we hypothesized that the slight differences between the spectra originating from these TCA probes could still enable multiplexing. To test this hypothesis, we treated A549 lung carcinoma epithelial cells with a mixture of d_7_-glc, d_3_-pyr, and d_5_-gln. We then acquired an hSRS image stack, tuning through several vibrational frequencies, and demonstrated robust unmixing performance with the least absolute shrinkage and selection operator^25^ (LASSO; **Fig. S3**), differentiating between the biomass originating from d_3_-pyr and d_5_-gln, and lipids and proteins derived from d_7_-glc (d_7_-glc_L_ and d_7_-glc_P_, respectively; **Fig. 2d**). The observed spatial distributions of each unmixed species correlate well with our single-probe SRS images (**Fig. 2a-b**), and with previous work on d_7_-glc and d_5_-gln, further validating these results.^20,21^ We achieved similarly robust unmixing in a mixture of d_7_-glc, d_3_-pyr, and d_3_-lactate (**Fig. S4**), demonstrating that these probes are broadly compatible in complex mixtures. MATRIX-SRS therefore allows direct comparison of TCA-derived biomass generation and spatial patterns between cell types for probing subcellular metabolic distribution.

### II. MATRIX-SRS reveals metabolic differences in epithelial-to-mesenchymal transition

We next applied MATRIX-SRS to investigate the TCA-related metabolic changes underlying EMT, which remains elusive at the metabolic level.^26^ During the EMT process, epithelial cells lose cell-cell adhesion and polarity while acquiring migratory and invasive mesenchymal properties,^27,28^ bearing deep relevance in understanding cancer progression and therapeutic resistance (**Fig. 3a**). Although recent studies have identified select metabolic alterations during EMT,^29^ most approaches rely on bulk measurements,^30^ transcriptional readouts,^31^ or other invasive methods^32^ that do not capture the spatiotemporal metabolic heterogeneity or rewiring at the subcellular level in living systems.

**Figure 3.**
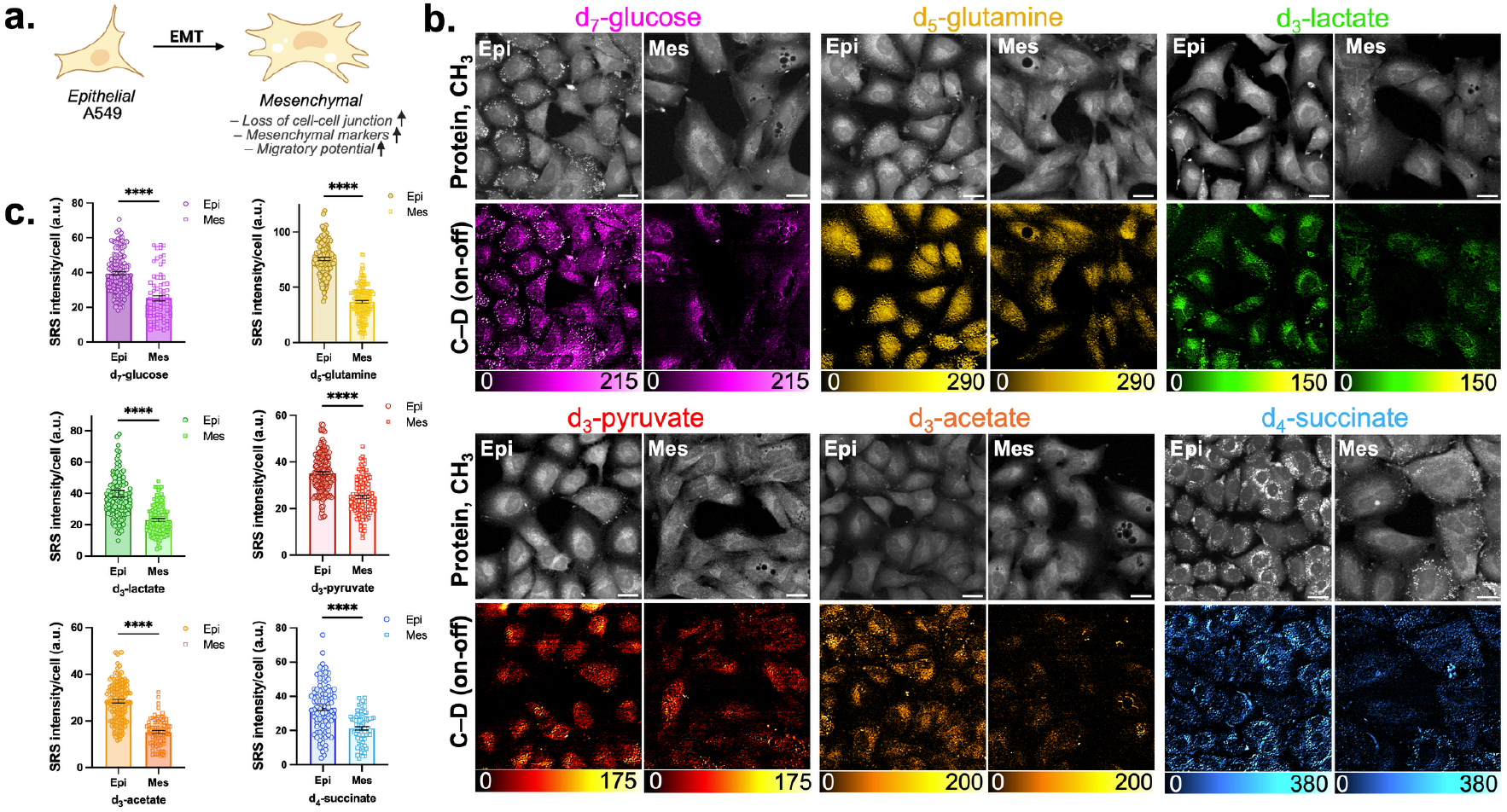
MATRIX-SRS for visualization of metabolic changes underlying EMT. **(a)** illustration showing epithelial-to-mesenchymal transition of A549 cells that is indicated by profound morphological changes, an increase in mesenchymal markers and migratory potential; **(b)** Live-cell SRS images targeted at the –CH_3_ channel (proteins, 2940 cm^−1^) and C–D (on-off) for epithelial (epi) and mesenchymal (mes) cells for the six metabolic probes: d_7_-glucose, d_5_-glutamine, d_3_-lactate, d_3_-acetate, d_3_-pyruvate, and d_4_-succinate. Scale bar: 20 µm; **(c)** Quantitative analysis of SRS signal of C–D bonds per cell. Data are presented as mean ± SEM from at least three independent experiments. Statistical significance was analyzed using two-tailed unpaired Student’s t-tests. ^****^p<0.0001.

To model EMT, we first induced A549 epithelial cells (epi) to adopt a mesenchymal (mes) phenotype using a commercial cocktail of proteins and antibodies, as confirmed by characteristic morphological changes such as cellular elongation (protein channel, **Fig. 3b**) and immunostaining (**Fig. S5**). Consistent with prior literature on MCF7 cells,^29^ metabolic imaging of d_7_-glc in live A549 cells showed reduced glucose metabolism in mesenchymal cells compared to epithelial cells (∼36%, **Fig. 3b-c**). Furthermore, MATRIX-SRS imaging revealed a global reduction in TCA-associated metabolic activity into the newly synthesized biomass during EMT, as indicated by the attenuated C–D signal for d_5_-gln (∼51%), d_3_-lactate (∼42%), d_3_-acetate (∼46%), d_3_-pyr (∼29%) and d_4_-succinate (∼36%) (**Fig. 3b-c**). These results highlight the potential of MATRIX-SRS in quantitatively investigating subcellular TCA-linked metabolism. In general, we observed a ∼30-40% reduction in total C–D signal in the biomass corresponding to the metabolic probes in EMT. Since many metabolic changes underlying EMT depend on the model system and remain controversial,^26,33^ we highlight the potential of MATRIX-SRS to better understand these longitudinal changes at the subcellular level in different systems.

### III. Complementary LC-MS SIL tracing provides molecular insight into reduced pyruvate utilization during EMT

To more accurately characterize the metabolic fate of the deuterium labeled probes and provide complementary confirmation of our SRS imaging results, we leveraged the high synergy between SRS microscopy and mass spectrometry.^22^ We chose d_3_-pyr, as a central intermediate in cellular metabolism uniquely positioned between the TCA cycle, glycolysis, protein synthesis, and DNL, for complementary *in vitro* tracing via LC-MS for EMT. We incubated the two cell types (epi and mes) using d_3_-pyr for 24 hours (**Fig. 4a**). As a metabolically active intermediate, pyruvate is known to anaplerotically incorporate into other small molecules through the TCA cycle. Given the enzymatic nature of these reactions, starting with d_3_-pyr should lead to the production of TCA intermediates (**Fig. 4b**) and many other biological molecules (**Fig. S6**) with highly specific deuterated motifs.

**Figure 4.**
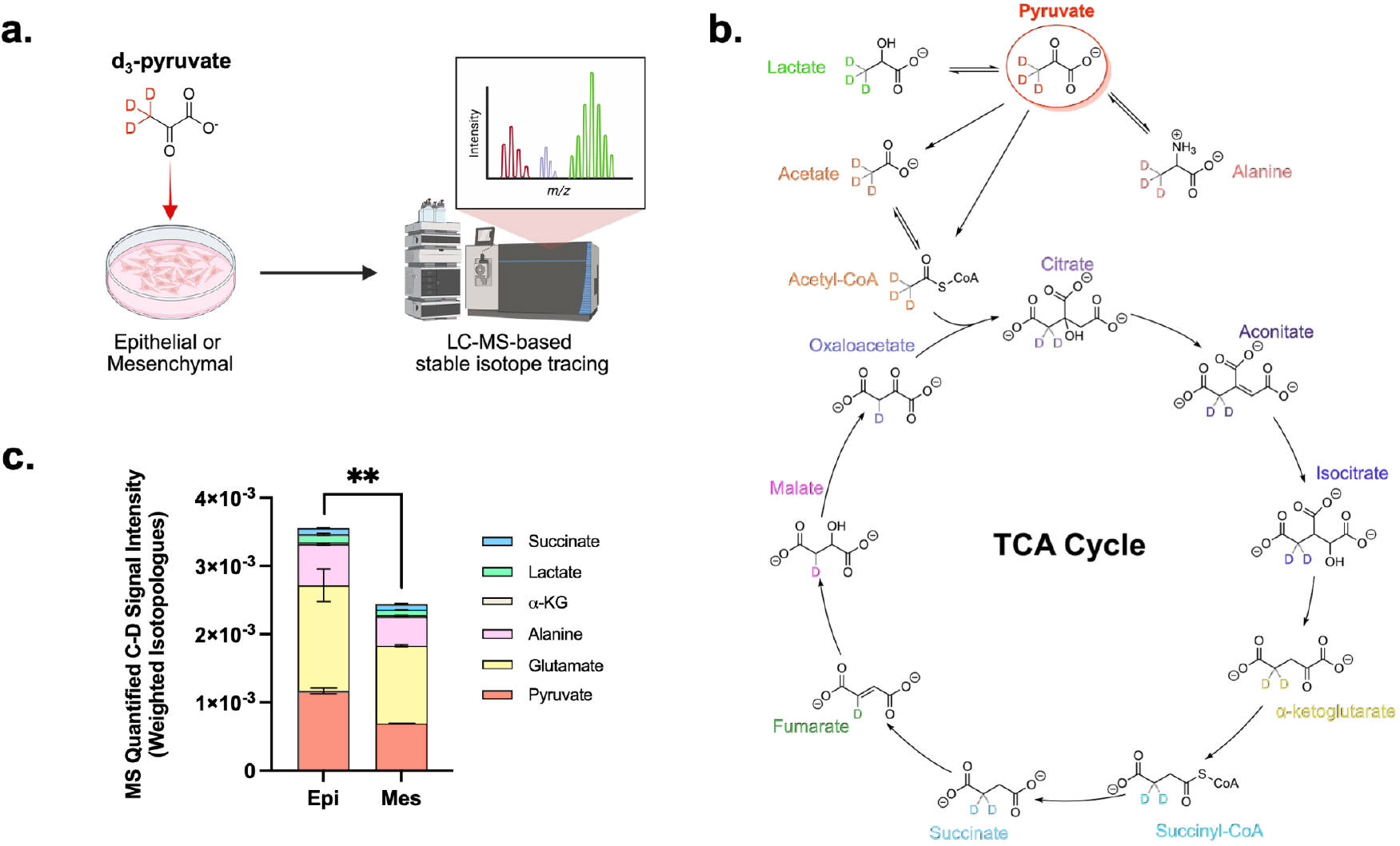
[3,3′,3″-^2^H_3_]-pyruvate *in vitro* tracing using LC-MS. **(a)** Illustration showing the *in vitro* tracing workflow using d_3_-pyruvate in A549 (epithelial or mesenchymal) cells; **(b)** Schematic of the flow of d_3_-pyruvate into the TCA cycle. **(c)** MS-quantified C–D incorporation (weighted isotopologues) of d_3_-pyr in TCA cycle intermediates. Data are presented as mean ± SEM from three independent experiments. Statistical significance was analyzed using two-tailed unpaired Welch’s t-test. ^**^p<0.01. Fractional isotopologue distributions are shown in **Fig. S7**.

Our *in vitro* LC-MS tracing data revealed that mesenchymal cells exhibit ∼30% lowered intracellular d_3_-pyr when compared to epithelial cells (**Fig. 4c**), showing excellent consistency with our SRS findings. More specifically, SIL tracing further indicated extensive conversion of d_3_-pyr into downstream metabolic species such as lactate, alanine, and citrate (**Fig. S7**), with the largest differences between epithelial and mesenchymal cells stemming from total glutamate and total pyruvate (**Fig. 4c**). Furthermore, examining the isotopologue distributions of individual metabolites, mesenchymal cells showed a significant attenuation of the M+3 and M+2 labeled fractions for acetylcarnitine (proxy for acetyl-CoA^33^) as well as citrate (M+1 and M+2 labeled fractions) which feeds into DNL,^34^ compared with their epithelial counterparts. Furthermore, pyruvate, which can be transaminated to alanine for protein synthesis,^35^ showed a reduced contribution to all alanine isotopologues in mesenchymal cells, indicating attenuated protein synthesis derived from d_3_-pyr. Taken together, these LC-MS tracing results complement our findings from MATRIX-SRS and provide deeper molecular specificity in understanding the metabolic fate of d_3_-pyr.

### IV. Hyperspectral MATRIX-SRS and DFT-derived spectral profiling

Inspired by the molecular specificity gained by LC-MS, we sought to develop a quantitative framework to determine concentration maps of metabolized C–D probes with hSRS. Upon close examination, we noted that the biomass spectra exhibited peaks at similar positions, differing primarily in their relative amplitudes (**Fig. 2c**). Such observations have been made previously with hSRS but remain poorly understood, with spectral assignments being largely empirical.^20^ Quantum mechanical calculations, such as DFT, have significant potential to aid in understanding experimental spectra, but these tools have not yet been applied to interpreting biomass spectra for two main reasons. First, DFT is scarcely used in SRS, due to the difficulty in determining an appropriate computational method and level of theory, the complexity of the target species (e.g., molecules with multiple possible protonation states) and a general perception that DFT is not quantitatively accurate for Raman spectroscopy.^34^ Second, to interpret *in situ* biomass spectra, computations must be performed at large scales, with the ability to calculate accurate spectra for dozens (if not hundreds) of diverse intermediate molecules, necessitating significant work in the implementation of DFT for metabolic SRS.

To address these challenges, we developed AutoDFT 2.0: a fully automated computational pipeline for calculating Raman spectra of selectively deuterated molecules requiring only ChemDraw structures as inputs (**Fig. S8**). AutoDFT 2.0 builds upon our previous work with automating DFT for calculations of quantitatively accurate vibrational spectra. In our previous work computing vibronic spectra of near-infrared organic dyes,^35^ we observed a significant improvement in computed spectral accuracy by using the solvation model with density (SMD) as an implicit solvent model.^36^ Compared with standard solvent continuum models, SMD is unique because it includes additional terms for short-range intermolecular interactions, such as acidity/basicity and surface tension, in addition to the dielectric description of most standard solvent models. Applying these same optimized methods, we calculated a theoretical Raman scattering spectrum for d_3_-pyr, obtaining excellent agreement with our experimental results (**Fig. 5a**). This high accuracy also generalizes to other molecules; we observed strong agreement across our TCA probes in frequency (R^2^ = 0.94; **Fig. 5b**) and similarly strong agreement in intensity (R^2^ = .93; **Fig. S9**). After performing detailed metabolic deuterium mapping (**Fig. S6**), we applied AutoDFT 2.0 to our entire library of mapped metabolites, totaling 69 unique structures (**Fig. S10**), generating comprehensive, physically informed knowledge about the vibrational features of myriad deuterium-labeled chemical structures.

**Figure 5.**
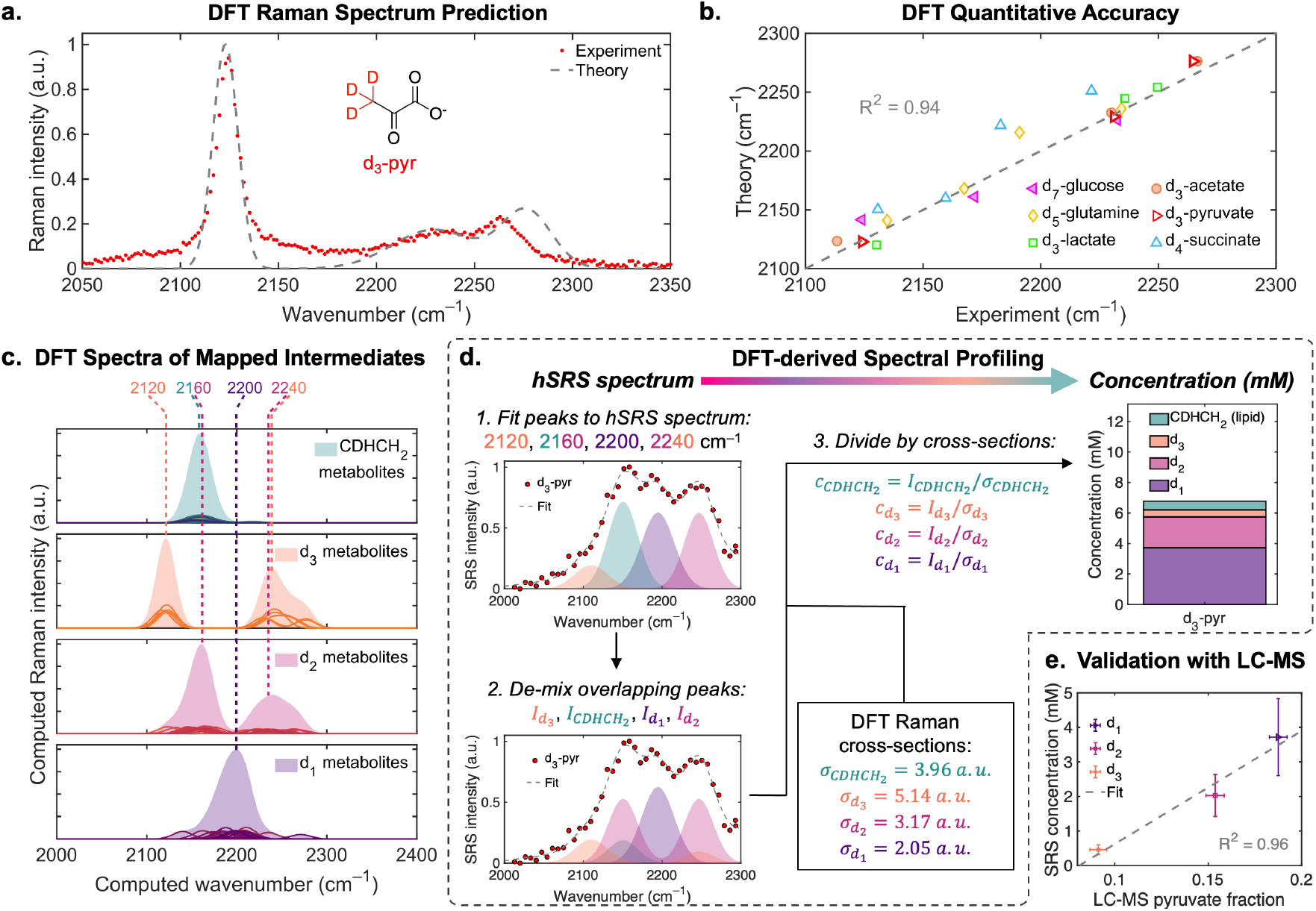
DFT spectral validation and compositional profiling. **(a)** Comparison of DFT-computed Raman spectrum for d_3_-pyr and an experimental Raman spectrum of d_3_-pyr in solution. **(b)** Comparison of calculated and experimental vibrational frequencies across all TCA probes in solution. **(c)** Aggregated DFT spectra of mapped deuterated TCA intermediates. Four distinct bands at 2120, 2160, 2200, and 2240 cm^−1^ are observed, characteristic of different labeled species. **(d)** DFT-derived spectral profiling. Fitting hSRS spectra at 2120, 2160, 2200, and 2240 cm^−1^ and dividing by DFT-derived cross-sections (**Fig. S12**) allows for *in situ* compositional profiling and absolute concentration determination for TCA metabolites. **(e)** Correlation between DFT-derived SRS concentrations and LC-MS-detected pyruvate fractions. Error bars are ± one standard deviation for LC-MS and ± 30% (estimated DFT error; see **Fig. S12**) for SRS.

Upon analyzing our DFT-computed spectra, we observed a striking clustering of spectral fingerprints based on the chemical structures of the deuterated molecules (**Fig. 5c** and **Fig. S10**). Singly deuterated intermediates (d_1_; **Fig. 5c** and **Fig. S10a**, purple) exhibit a local C–D stretching mode at ∼2200 cm^−1^. For molecules with deuterated methylenes (d_2_; **Fig. 5c** and **Fig. S10b**, pink), the C–D stretching band is split into a redshifted, symmetric stretching band (∼2160 cm^−1^) and a blueshifted, asymmetric stretching band (∼2240 cm^−1^). Fully deuterated methyl groups (d_3_; **Fig. 5c** and **Fig. S10c**, orange) exhibit a further redshifted symmetric stretching band (∼2120 cm^−1^) and two tightly clustered asymmetric stretching bands (both ∼2240 cm^−1^). Computed spectra for deuterated fatty acids (of the form –(CDHCH_2_)_n_– from our reaction mapping; **Fig. S6**) exhibit a single strong band at ∼2160 cm^−1^ (**Fig. 5c** and **Fig. S10d**, teal). Furthermore, the fatty acid scattering intensity is highly correlated with the number of deuteriums in the chain (R^2^ = 0.93; **Fig. S11**), allowing us to readily quantify lipid signals on a per-deuterium basis (CDHCH_2_).

To convert from hSRS spectra to concentration, we leveraged that SRS signal is proportional to the product of the concentration of an analyte (within the focal volume) and the scattering cross-section of the analyte. With the consistency achieved from DFT computed intensities with experimental data, we further derived average scattering cross-sections for each of the four deuterated motifs (d_1_, d_2_, d_3_, and CDHCH_2_), which we divide by our experimentally measured intensities to determine concentrations (**Fig. 5d**; see also **Fig. S12**). Applying this framework for d_3_-pyr, we obtained a total deuterated biomass concentration of 6.8±2.0 mM. On a per-species basis, we find that most of the deuterium biomass is found in d_1_ species (∼4 mM) with progressively less in d_2_ and d_3_ species (**Fig. 5d**; see also **Table S1**). Relatively little deuterium biomass (<1 mM) is found in the internal positions in aliphatic lipid tails (CDHCH_2_), but an appreciable amount of the measured d_1_, d_2_, and d_3_ biomass may still originate from lipids due to head group functionalization and the terminal methyl groups of fatty acids, with the remainder most likely originating from proteins (**Fig. S6**). To our knowledge, this is the first time that absolute concentrations of metabolized deuterium probes have been determined *in situ* by SRS microscopy.

In principle, our SRS quantification results should be comparable with our LC-MS-quantified isotopologue distributions, where the found M+1 peak corresponds to d_1_ species, M+2 corresponds to d_2_, and M+3 corresponds to d_3_. Upon attempting this comparison, we found surprisingly excellent agreement (R^2^ = 0.96; **Fig. 5e**), with LC-MS similarly indicating globally lower abundances for more highly deuterated species (**Fig. S7**). These results validate our DFT-derived hSRS spectral profiling as a means of determining absolute *in situ* concentrations of metabolized C–D probes with quantitative accuracy. We also applied this quantitative pipeline to d_7_-glc, a much more well-studied Raman metabolic probe, allowing for direct quantification of the compositions of d_7_-glc_L_ and d_7_-glc_P_ (**Fig. S13**).

Biochemically speaking, the conversion of our starting material, d_3_-pyr, into d_2_ and d_1_ species indicates the active metabolism of d_3_-pyr (where unreacted d_3_-pyr would exhibit purely d_3_ biomass). For example, the conversion of d_3_-pyr into d_2_-citrate (through acetyl-CoA; **Fig. 4b**) necessitates a deuterium abstraction, leading to the loss of d_3_ biomass and gain of d_2_ biomass. Similarly, the dehydrogenation of d_2_-succinate to produce d_1_-fumarate converts d_2_ biomass into d_1_ biomass. These TCA intermediates can then be subsequently incorporated into stable downstream proteins or lipids, such as the feeding of citrate into DNL or conversion into amino acids, leading to build-up of deuterated biomass from d_3_-pyr turnover.

For the first time, MATRIX-SRS allows for live-cell-compatible quantification of deuterium biomass. In tandem with LC-MS to obtain metabolite-specific isotopologue distributions, our integrative approach provides a uniquely detailed perspective of global cellular metabolism. Moreover, the enrichment of different C–D species could imply the preference of one metabolic pathway over another for a given metabolic precursor. For example, observing appreciable d_3_ biomass in newly synthesized proteins could indicate rapid conversion of d_3_-pyr into d_3_-alanine via transamination (**Fig. 4c**), while conversion into other amino acids generally entails deuterium abstraction (**Fig. S6**), meaning d_1_ and d_2_ species are expected to be the dominant sources of total protein biomass (as with d_7_-glc_P_; **Fig. S13**). Moreover, the conversion of d_3_-pyr into d_3_-lactate (mediated by NADH) can report the redox state of the cell. Our approach could hence facilitate quantitative comparisons between different treatment conditions to investigate the intricate metabolic network.

### V. Pharmacological perturbation of d_3_-pyruvate metabolism

To quantitatively investigate metabolic perturbation, we next pharmacologically treated these metabolic pathways to visualize their outcomes using SRS imaging in live A549 cells incubated with d_3_-pyr. We first inhibited ATP-citrate lyase (ACLY), the enzyme that converts cytosolic citrate into acetyl-CoA (and subsequently fatty acids) following a series of biochemical reactions in DNL^37^ (**Fig. 6a**). Surprisingly, pharmacological perturbation of ACLY using three distinct small molecule inhibitors (SMIs) – bempedoic acid (BA, or ETC-1002)^38^, SB-204490^39^, and BMS-303141^40^ – did not significantly attenuate the C–D signal from d_3_-pyr in live A549 cells when compared to a control group of untreated cells (**Fig. S14**). These results are consistent with previous findings suggesting that pyruvate may be converted into acetate and then into acetyl-CoA through ACLY-independent pathways.^41,42^

**Figure 6.**
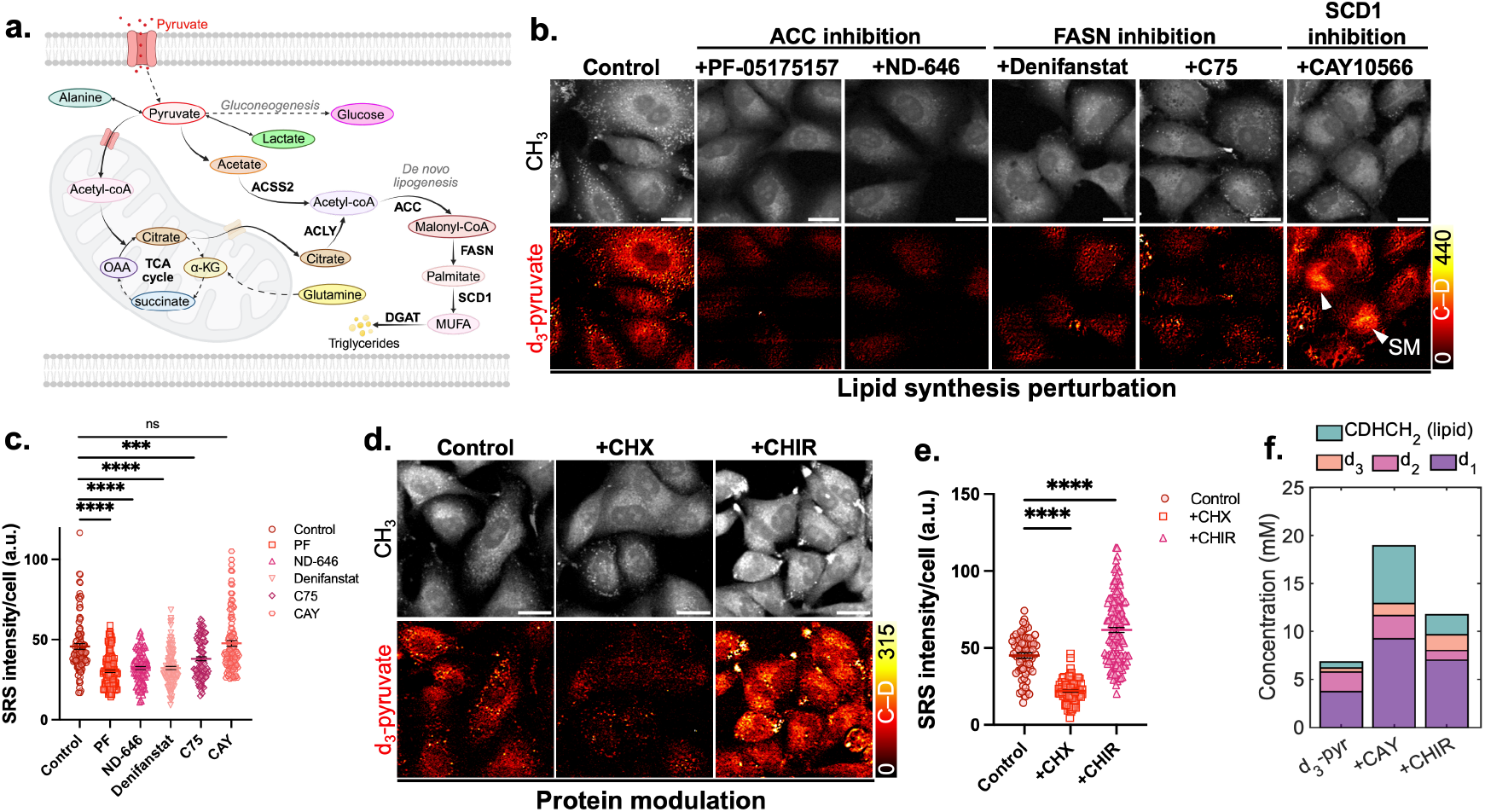
Pharmacological intervention of pyruvate metabolism. **(a)** Schematic of pyruvate metabolism in cells leading to *de novo* lipogenesis (DNL). ACC = acetyl-CoA carboxylase, ACLY = ATP-citrate lyase, ACSS2 = acyl-CoA synthetase short-chain family member 2, FASN = fatty acid synthase, SCD1 = stearoyl-CoA desaturase 1, DGAT = diacylglycerol acyltransferase, MUFA = monounsaturated fatty acid, SFA = saturated fatty acid; **(b)** Live-cell SRS images of A549 cells incubated with 30 mM d_3_-pyruvate for 48 hours targeted at the protein channel (–CH_3_, 2940 cm^−1^) and C–D (on-off) for (i) control (no drugs), (ii) ACC inhibition (using 1 µM PF-05175157, or 1 µM ND-646), (iii) FASN inhibition (using 10 µM denifanstat, or 75 µM C75), or (iv) SCD1 inhibition (using 1 µM CAY10566) that resulted in SFA-induced solid membrane (SM); **(c)** Quantification of C– D intensity per cell upon lipid metabolism perturbation using SMIs (from b); **(d)** Live-cell SRS images of A549 cells incubated with 30 mM d_3_-pyruvate for 48 hours targeted at the protein channel (–CH_3_, 2940 cm^−1^) and C–D (on-off) for (i) control (no drugs), (ii) 1 µM cycloheximide, or (iii) 3.5 µM CHIR-99021 inhibition; **(e)** Quantification of C–D intensity per cell upon protein metabolism perturbation using SMIs (from d). Data are presented as mean ± SEM from at least three independent experiments. Statistical significance was analyzed using two-tailed unpaired Student’s t-tests. ns = no statistical difference, ^***^p<0.001, ^****^p<0.0001; **(f)** DFT-derived quantification of d_3_-pyr-derived biomass (untreated), SFA-enriched SM domains (+CAY), and cytoplasmic d_3_-pyr-derived biomass following CHIR treatment. All scale bars: 20 µm.

We next perturbed acetyl-CoA carboxylase (ACC), the enzyme that mediates fatty acid synthesis by converting acetyl-CoA into malonyl-CoA, using two specific SMIs (PF05175157^43^ and ND-646^44^). SRS imaging showed a dramatic decrease in d_3_-pyr signal in A549 cells treated with these inhibitors compared to the control (**Fig. 6b-c**). Furthermore, inhibition of fatty acid synthase (FASN), which catalyzes the synthesis of palmitate from acetyl-CoA and malonyl-CoA,^37^ using denifanstat^45^ and C75^46^ also resulted in decreased d_3_-pyr signal. These results indeed suggest that pyruvate metabolizes through ACC and FASN enzyme activity. Lastly, inhibition of stearoyl-CoA desaturase 1 (SCD1), which converts saturated fatty acids (SFAs) to monounsaturated fatty acids (MUFAs), using CAY10566, elicited an altered response; we found that the C– D intensity in these cells remained comparable to control cells (**Fig. 6b-c**). Interestingly, these cells exhibited enriched C–D signal in cytoplasmic pools, which we classify as solid membrane (SM) domains, as discovered previously in mesenchymal melanoma cancer cells.^47^ The SM formation results from SCD1-induced imbalance between saturated and unsaturated fatty acids (due to increased SFAs) and is characterized by its distinct Raman spectrum (**Fig. S15a**). Taken together, our findings corroborate that a significant portion of d_3_-pyr gets incorporated into downstream lipids via DNL.

Since d_3_-pyr is also expected to contribute to newly synthesized proteins in our results, we next modulated protein synthesis by either inhibiting it using cycloheximide (CHX)^48^ or promoting it using CHIR-99021 (CHIR) through inhibition of glycogen synthase kinase 3 (GSK-3).^49^ As expected, inhibition of protein synthesis with CHX significantly decreased the C–D signal in cells, whereas upregulation of protein synthesis with CHIR increased the overall C–D intensity (**Fig. 6d-e**). Additionally, hSRS imaging revealed an increase in the ∼2200 cm^−1^ and ∼2240 cm^−1^ bands in cells treated with CHIR (**Fig. S15b**), which are attributed to d_1_- and (primarily) d_2_-labeled metabolites including proteins, in contrast to cells that primarily exhibited lipid-rich subcellular SM pools (SCD1 inhibition in **Fig. 6b, Fig. S15a**) attributed to lipid (CDHCH_2_) metabolites.

We further quantified the deuterium distribution in these SFA-rich SM domains (**Fig. S15a**) as well as in the cytoplasm (**Fig. S15b**) of CHIR-treated A549 cells using our DFT-derived MATRIX-SRS workflow. In SM domains (**Fig. 6f**, +CAY), we observed a tenfold increase in deuterium biomass in lipids (6.0±1.8 mM, CDHCH_2_, teal) and appreciable increases in d_1_ (purple) and d_3_ (orange) metabolite concentrations compared with the d_3_-pyr control (**Fig. 6f**, d_3_-pyr). The increase in d_3_ (orange) concentration is especially interesting and may indicate faster conversion of d_3_-pyr into d_3_-acetyl-CoA, and subsequent incorporation into the terminal methyl groups in fatty acids (**Fig. S6**). On the other hand, in cells treated with CHIR (**Fig. 6f**, +CHIR), we found increased concentrations of d_1_ (purple), d_3_ (orange, most likely from the conversion of d_3_-pyr to d_3_-alanine; **Fig. 4b**), and lipid (CDHCH_2_, teal) deuterium, but a surprising reduction in d_2_ biomass (pink) compared with the d_3_-pyr control (**Fig. 6f**, d_3_-pyr). We reason that the reduction in the d_2_/d_1_ ratio may be an indicator of overall higher TCA cycle activity, where the progression of the cycle naturally leads to deuterium abstraction when proceeding through fumarate (**Fig. 4b**). Therefore, the decreased relative d_2_ abundance observed upon treatment with CHIR is consistent with broadly upregulated metabolic activity. In this manner, MATRIX-SRS holds significant potential as a live-cell-compatible method to investigate the impacts of drugs on metabolism by directly visualizing and quantifying metabolome-wide changes in absolute concentrations.

## Discussion

Targeted SRS microscopy using small vibrational tags enables visualization of dynamic metabolic turnover and the incorporation of labeled substrates into distinct cellular biomacromolecules.^11,16^ However, the inherent complexity, rapid dynamics and low abundance of intermediate metabolites connecting the initial probe to the final biomass limit our ability to fully map the metabolic atlas. This highlights the need for expanding the repertoire of probes capable of visualizing these transient metabolic species, and a robust and quantitative analysis platform with generally comparable units (e.g., absolute concentration) across multiple experiments and model systems.

We demonstrate the broad applicability of MATRIX-SRS in visualizing both TCA-related anabolism and catabolism in live HeLa cells and neuronal co-cultures. We then applied this methodology to investigate the metabolic reprogramming underlying EMT in live A549 adenocarcinoma cells. MATRIX-SRS revealed a ∼30-40% global attenuation of TCA-associated metabolic incorporation of d_7_-glucose, d_5_-glutamine, d_3_-lactate, d_3_-acetate, d_3_-pyruvate and d_4_-succinate in mesenchymal cells compared to their epithelial counterparts. Importantly, we established a physics-informed MATRIX-SRS quantification, employing high-throughput density functional theory calculations to obtain absolute concentrations of metabolized deuterium biomass based on hSRS spectra.

We focused on pyruvate, which occupies a central position between sugar processing, the TCA cycle, amino acid synthesis, and lipid synthesis (**Fig. S6**), to further dissect its metabolic role. We first complemented our SRS imaging results with SIL tracing via LC-MS, which confirmed a ∼30-40% downregulation of pyruvate utilization on a per-metabolite basis in mesenchymal cells and demonstrating the high synergy between LC-MS and SRS. We further applied quantitative MATRIX-SRS analysis to d_3_-pyr, achieving excellent consistency with SIL tracing, granting previously inaccessible pathway-dependent insights derived from precise biomass quantification.

SRS intensity depends on both the abundance and the scattering cross-section of the analyte molecule. For metabolically stable probes (e.g., alkyne-tagged molecules like 5-ethynyl-2’-deoxyuridine, EdU), absolute concentrations can be determined *in situ* by generating a calibration curve in solution. However, for metabolically active probes that undergo metabolic turnover (e.g., deuterium-labeled metabolites, as with our TCA probes), the observed biomass comprises metabolites of both unknown concentrations and unknown cross-sections, resulting in an underdetermined system. These downstream products are highly heterogeneous, with hundreds of possible structures of varying levels of deuteration (**Fig. S10e**). Moreover, solution reference spectra are not readily acquired for these species, which are transient, low-abundance, and immensely challenging to selectively isolate or prepare synthetically on large scales. As such, SRS on metabolically active probes has been largely qualitative in intensity, instead focusing on spatial heterogeneity or unmixing from empirically derived spectral differences.^20^

In principle, computational methods have long been capable of complementing experimental data to provide cross-section estimates for transient species. Compared with empirical modeling, physics-based calculations are derived from an underlying, internally consistent set of fundamental principles, facilitating much deeper insights and broader generalization and prediction. However, the application of DFT calculations for Raman spectra of small molecule metabolites faces several barriers, including method selection, validation, and application to dozens of diverse metabolites (effectively necessitating automation), leading to largely empirical spectral assignments and limited interpretability. ^34^

Our work shows that careful optimization of the computational methods (including the choice of an accurate but cost-effective solvent model^36^) can allow for quantitatively accurate computations of both Raman scattering frequencies and intensities (**Fig. 5b** and **Fig. S9**). Based on these findings, we applied DFT calculations at large scale to nearly 70 TCA-linked metabolites (**Fig. S10**) and derived cross-section estimates for four spectrally distinct metabolite classes (d_1_, d_2_, d_3_, and CDHCH_2_), allowing for *in situ* quantification of metabolically active deuterium probes in absolute concentrations with SRS for the first time. Moreover, we expect these quantification methods and metabolite classes to generalize broadly to other metabolically active deuterium probes, since these motifs are highly conserved through the metabolic reaction network surrounding the TCA cycle (**Fig. S6**).

Encouragingly, we observe remarkably good agreement between the LC-MS M+1/M+2/M+3 pyruvate fractions and our SRS-derived d_1_/d_2_/d_3_ concentrations (**Fig. 5e**), laying a foundation for future *in situ* metabolomic quantification with MATRIX-SRS. A unique strength of SRS-based spatial metabolomics is the potential for dynamic live-cell imaging, where subcellular metabolism could be observed directly by watching the evolution of the d_1_/d_2_/d_3_/CDHCH_2_ distribution over time.

We should also note a few key limitations. In our current work, we employ slightly elevated concentrations of our TCA probes to achieve robust C–D signals with high signal-to-noise ratios, although the concentrations used (e.g., 10-30 mM for pyruvate) are still lower than those typically required for other techniques, such as MRI (∼69 mM).^50^ We carefully characterized cell states following probe incubation and observed no obvious toxicity effects. Importantly, our methods (tandem SRS and LC-MS, informed by physics-based modeling) are broadly generalizable to Raman metabolic imaging, and continued technical developments^51^ may grant much improved sensitivity to allow investigations at more physiologically relevant concentrations (such as ∼5.8 mM for pyruvate^23^). Additionally, our current study focuses on late-time endpoints to study bulk metabolism (and allow sufficient time to overcome deuterium kinetic isotope effects), while the dynamics of the TCA cycle remain relatively underexplored in metabolic imaging. Lastly, our DFT-derived quantification of deuterated biomass currently only allows for bulk classification into four classes of metabolites (d_1_, d_2_, d_3_, and CDHCH_2_). Although we complement our SRS imaging with LC-MS SIL tracing to gain deeper molecular specificity, an ideal live-cell spatial metabolomics approach would grant both subcellular localization and per-molecule identification from imaging alone. Multimodal imaging modalities,^52^ imaging techniques based on higher-dimensional spectroscopies,^53^ and new approaches integrating SRS and machine learning have significant potential to achieve greater molecular-level detail towards this ideal.

In the future, we also envision expanding MATRIX-SRS to other metabolic probes, such as fumarate, malate, and oxaloacetate, in living systems. Additionally, we expect that deeper functional investigations of our emerging TCA probes, as we have done here with d_3_-pyr, will also prove highly informative; for example, detailed study of d_3_-lactate could yield fascinating insights into the lactate shuttle hypothesis. By harnessing characteristic Raman spectral features and incorporating novel metabolic probes, we envision quantitative spatial metabolomics for longitudinal analyses. MATRIX-SRS, incorporating both high-resolution spectro-microscopy and DFT calculations, provides a uniquely quantitative basis for far-field imaging of metabolism with minimally perturbative probes, complementing existing strategies for metabolic profiling of living systems.

## Materials & Methods

### Cell Culture and Metabolic Labeling

HeLa cells (ATCC, CCL-2), and A549 cells (gift from the Elowitz lab, Caltech) were each cultured in DMEM (Gibco, Cat. #11965092) supplemented with 10% FBS (Corning, Cat. #35-015-CV) and 1× antibiotic-antimycotic (Gibco, Cat. #15240062) at 37 °C with 5% CO_2_. To induce EMT, A549 cells were seeded and cultured in DMEM containing 1x StemXVivo EMT-inducing supplement (R&D systems, Cat. #CCM017) for 5 days, with media refreshed every 2-3 days.

For neuronal co-culture, primary hippocampal neurons were prepared from neonatal (P0-P1) Sprague-Dawley rat pups (CD (Sprague-Dawley), IGS rat, Charles River) in accordance with an IACUC-approved protocol (IA22-1835). Following dissection, mice brains were transferred into ice-cold Hank’s balanced salt solution (Gibco, Cat. #14-025-092). Hippocampi were isolated, cut into ∼0.5 mm fragments, and digested with 5 mL of 0.25% Trypsin-EDTA (Gibco, Cat. #25200056) at 37 °C and 5% CO_2_ for 20 min. Digestion was quenched with 5 mL of DMEM containing 10% FBS. The tissue was transferred into 2 mL of neuronal culture medium (Neurobasal A supplemented with B-27, 2 mM GlutaMAX, and 1x penicillin-streptomycin; Thermo Fisher), then pelleted by brief centrifugation and resuspended in neuronal culture medium. Cells were dissociated by gentle trituration and diluted to a final density of 1×10^5^ cells/mL. For plating, 0.7 mL of the cell suspension was added per well of a 24-well plate containing 12 mm circular coverslips pre-coated with poly-D-lysine and laminin. Coated coverslips were prepared in advance by incubating sterile coverslips with 100 µg mL^-1^ poly-D-lysine (Sigma) at 37 °C for 24 h, rinsing twice with ddH_2_O, and subsequently coating with 10 µg/mL laminin (Gibco) overnight under the same conditions. Cell cultures were maintained at 37 °C and 5% CO_2_, and half of the medium was replaced every five days with a pre-warmed neuronal medium. The metabolic probes were added on ∼DIV11 at their respective concentrations and for the specified duration of time.

The metabolic probes used for SRS imaging were 5 mM 2,3,3,4,4-d_5_-L-glutamine (Cambridge Isotope Laboratories, Cat. #DLM-1826), 30 mM 3,3,3-d_3_-sodium L-lactate (Cambridge Isotope Laboratories, Cat. #DLM-9071), 10-30 mM d_3_-sodium pyruvate (Cambridge Isotope Laboratories, Cat. #DLM-6068), 30 mM d_3_-sodium acetate (Cambridge Isotope Laboratories, Cat. #DLM-3126) and 30-50 mM d_4_-succinic acid, disodium salt (Cambridge Isotope Laboratories, Cat. #DLM-2307) and 25 mM 1,2,3,4,5,6,6-d_7_-D-glucose (Cambridge Isotope Laboratories, Cat. #DLM-2062) in cell media containing no unlabeled counterparts of the respective metabolic probes for the designated time points.

For metabolic imaging of EMT, deuterium-labeled probes were added 3-4 days after cell seeding (for either epithelial or mesenchymal cells) on cover slips, at approximately ∼50-60% cell confluency. The concentrations and incubation time points of the metabolic probes were adjusted to 25 mM d_7_-glucose for 24 hours, 2 mM for d_5_-glutamine for 24 hours, 20 mM for d_3_-pyruvate for 24 hours, 30 mM d_3_-acetate for 48 hours, 30 mM d_3_-lactate for 48 hours, and 30 mM d_4_-succinate for 5 days in cell media containing no unlabeled counterparts of the corresponding probes.

### Drug Perturbation of Lipid and Protein Metabolism

For pharmacological perturbation of lipid metabolism, A549 cells were treated with 10 µM ETC-1002 (MedChemExpress, Cat. #HY-12357), 20 µM SB-204490 (MedChemExpress, Cat. #HY-16450), 2 µM BMS-303141 (MedChemExpress, Cat. #16107), 75 µM C75 (MedChemExpress, Cat. #HY-12364), 10 µM Denifanstat (MedChemExpress, Cat. #HY-112829), 1 µM PF-05175157 (MedChemExpress, Cat. #HY-12942), 1 µM ND-646 (MedChemExpress, Cat. #HY-101842), or 1 µM CAY10566 (MedChemExpress, Cat. #HY-15823) for 48 hr concurrently with 30 mM d_3_-pyruvate labeling.

For protein modulation experiments, A549 cells were treated with 1 µM cycloheximide (CHX, Sigma-Aldrich, Cat. #C7698-1G) or 3.5 µM CHIR99021 (MedChemExpress, Cat. #HY-10182) for 48 hr with concurrent d_3_-pyruvate labeling.

### SRS Microscopy

The SRS microscopy system was configured as described previously.^54^ Briefly, images were acquired with an 80 µs pixel dwell time to achieve an image acquisition speed of 8.52 s per frame for a 320×320-pixel field of view (FOV) at a 0.497 µm/pixel resolution. The pump beam wavelength was set to 791.3 nm for the protein (–CH_3_ channel, 2940 cm^−1^), 797.3 nm (–CH_2_ channel, 2845 cm^−1^), 842.9 nm (C–D channel for d_5_-glutamine, 2167 cm^−1^), 844.0 nm (C–D channel for d_7_-glucose/d_3_-pyruvate/d_3_-acetate/d_4_-succinate/d_4_-citrate, 2150 cm^−1^), 845.6 nm (C–D channel for d_3_-lactate, 2129 cm^−1^) and 855.0 nm (off-resonance, 2000 cm^−1^).

For hSRS, the wavelength of the pump laser was tuned from 834.0 nm to 855.0 nm with a step size of 0.5 nm. The hSRS spectrum for d_5_-gln was reproduced from a previous report.^22^ All images were processed and color-coded using ImageJ software.

For C–D quantification, on-resonance images were processed by subtracting the corresponding off-resonance images to result in the C–D (on-off) images. C–D (on-off) images were further processed with the Fiji ‘nonuniform background removal’ plugin before performing a uniform background subtraction on each image. The protein image was used to generate a mask for each cell utilizing the ‘IdentifyPrimaryObjects’ module in CellProfiler 4.2.8^55^ with the ‘Minimum Cross-Entropy’ thresholding method. The mask was then used as a reference to quantify the final processed C–D image utilizing the ‘MeasureObjectIntensity’ module and exported to Excel using the ‘ExportToSpreadsheet’ module. MS Excel, GraphPad Prism, and MATLAB (R2024b, MathWorks) were used for data processing and plotting.

### Spontaneous Raman Microscopy

Spontaneous Raman spectra of analytes were acquired using a Horiba Xplora plus confocal Raman spectrometer equipped with a 532 nm YAG laser (12 mW) and a 100x, 0.9 N.A. objective (MPLAN N, Olympus) with a 500 µm hole and a 100 µm slit. Data were collected with a 10 s integration time and averaged over 10 accumulations using LabSpec6 software. Background signals from solvents were subtracted from the analyte spectra, followed by baseline correction and normalization.

### Immunofluorescence

Immunostaining of vimentin in epithelial and mesenchymal cells was performed as described previously.^29^ Cell nuclei were labeled with DAPI (Thermo Fisher, Cat #. D1306) and imaged using an excitation of 358 nm and an emission peak around 461 nm. Correlative SRS images were acquired at the protein channel (2940 cm^−1^) as described above using the same microscope.

### Multiplex Image Unmixing

Multiplex images were unmixed using the least absolute shrinkage and selection operator (LASSO) as previously described (**Supplementary Figure 6**). hSRS stacks were pre-processed by non-local means denoising^56^ and non-uniform background removal.^57^ After pre-processing, hSRS stacks were power-corrected and background-corrected, then unmixed via LASSO (λ = 0.3) on a per-pixel basis.

### DFT Calculations

Density functional theory (DFT) calculations were carried out using our AutoDFT computational pipeline (https://github.com/pkocheril/AutoDFT). ChemDraw (v23.1.2) was used to prepare structure files for each metabolite of interest. Through AutoDFT 2.0, we generated 3D geometries with selective deuteration and performed a classical energy minimization with OpenBabel.^58^ We then automatically generated and submitted job files for Gaussian 16 (rev. B.01, Gaussian Inc.)^59^, performing a coarse geometry optimization (HF/STO-3G), followed by fine optimization and harmonic frequency calculations (B3LYP/6-31G(d,p)) with implicit solvation in dimethyl sulfoxide using the solvation model with density (SMD).^36^ Natural bond orbital analysis was employed and symmetry was disabled. Computed normal mode frequencies were scaled by 0.9658. Raman spectra were calculated by applying Gaussian broadening (25 cm^−1^ full-width at half-maximum; comparable to biomass spectral linewidths) to each of the calculated peaks. GaussView (v6.1.1) was used for additional visualization.

### DFT-derived SRS Quantification

SRS intensity can be modeled as *SRS* ∝ *c* · *σ* · *I*_*Pump*_ · *I*_*Stokes*_, where *c* is the concentration of scattering molecules within the focal volume, *σ* is the scattering cross-section, and *I*_Pump_ and *I*_Stokes_, are the powers of the pump and Stokes beams, respectively. Thus, we can relate concentration to SRS intensity as 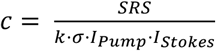, where k is a constant of proportionality encompassing various experimental factors such as scattering collection efficiency and photodiode sensitivity. Here, we use DFT to calculate the mean scattering cross-sections across our four classes of deuterated metabolites (d_1_, d_2_, d_3_, and lipids (CDHCH_2_)), calculated by summing the total computed intensity and dividing by the number of metabolites in the class (in a.u.). To obtain *k*, we measured an SRS spectrum of a 1 M aqueous solution d_3_-pyr (for our system, *k* = 0.12 mW^−2^ M^−1^ (a.u.)^−1^), thus allowing us to convert between SRS intensity and molar concentration (**Fig. S12**).

### [3,3′,3″-^2^H_3_]-pyruvate SIL Tracing Using LC-MS

A549 cells were cultured in DMEM containing 10% dialyzed FBS (Gibco, Cat.# A3382001) on 6-cm cell culture dishes. After EMT induction, epi and mes cells were treated with 10 mM d_3_-pyr. Following 24 hours, cell metabolism was quenched by rapidly aspirating the medium, rinsing with 150 mM NH_4_OAc (pH adjusted to 7.3, Fisher Scientific, Cat. #ICN19384880) and adding 80% ice-cold methanol containing 1 µm norvaline (Millipore Sigma, Cat. #53721-100MG) as the internal standard. Samples were incubated at -80 °C for 30 min, scraped into low-bind tubes and centrifuged at maximum speed for 30 min. Supernatants were stored at -80 °C until LC-MS analysis. Prior to injection, extracts were thawed on ice, centrifuged for 10 min, and clarified for LC-MS.

Metabolites were separated using a Waters BEH z-HILIC column (150 x 2.1 mm, 2.7 µm) connected to a Thermo Scientific™ Orbitrap Exploris™ 480 mass spectrometer with a heated electrospray ionization (H-ESI) source. The column temperature was maintained at 25° C, and the autosampler at 4° C. The mobile phase consisted of Solvent A: 10 mM ammonium bicarbonate in water (pH ∼9.1) and Solvent B: 95% acetonitrile/5% water. Chromatographic separation was conducted at 0.18 mL/min with the following gradient: 0-1 min, 95% B; 1-15 min, linear decrease to 45% B; 15-19 min, 45% B; 19-22 min, ramp back to 95% B; 22-34.5 min, 95% B for re-equilibration.

MS data were acquired in data-dependent acquisition (DDA) mode with fast polarity switching. Full MS^1^ scans were collected at 120,000 resolution (m/z 70-1000) with an AGC target of 1 x 10^6^. MS/MS scans were acquired using a TOP2 ddMS^2^ method with higher-energy collisional dissociation (HCD) at stepped collision energies of 10% and 40%, resolution 30,000, and isolation window 0.7 m/z. Dynamic exclusion, isotope exclusion, monoisotopic precursor selection, and mild ion trapping were enabled to enhance spectral quality.

Raw LC-MS data were converted to mzXML format using MSConvert and processed in El MAVEN (v0.12.0). Peak detection, chromatogram alignment, and automated isotope annotation were performed with default settings. Retention times were normalized using external standards, ion intensities corrected for matrix effects, and peaks manually curated as needed. Processed peak tables were exported in CSV format for analysis of isotopologue distributions.

To account for the contribution of each labeling state, isotopologues were weighted by the number of incorporated labeled atoms (e.g., M+1 by 1, M+2 by 2, M+3 by 3, etc.). This weighting demonstrates the relative contribution of each isotopologue to the overall C–D signal. These weighted values were then multiplied by previously published metabolite concentrations^23^ and summed to determine the total metabolite C–D burden. This approach ensures that mass spectrometric quantification accurately reflects the extent of deuterium incorporation, mirroring the Raman-derived signal. Final data were expressed either as the fraction of the total signal from labeled species or as relative isotopologue abundances, depending on the downstream analysis.

## Data Availability

The raw data supporting this study are available upon reasonable request from the author. The code for AutoDFT is publicly available on GitHub (https://github.com/pkocheril/AutoDFT).

## Acknowledgements

R.S.C. acknowledges support from the Biotechnology Leadership Pre-doctoral Training Program (BLP) in the Donna and Benjamin M. Rosen Bioengineering Center at Caltech. P.A.K. and A.C. are grateful for financial support from a National Science Foundation Graduate Research Fellowship (DGE-1745301 and DGE-2139433, respectively). P.A.K. is additionally grateful for financial support from a Hertz Fellowship. The computations presented here were conducted with the Resnick High Performance Computing Center, a facility supported by the Resnick Sustainability Institute at the California Institute of Technology. L.W. is a Heritage Principal Investigator supported by the Heritage Medical Research Institute and also acknowledges support from a CZI dynamic imaging grant. Select figures were created using biorender.com. We thank Dr. Ryan Leighton, Dr. Jacob Kirsh, Jacob-Parres Gold, and Noor Naji for helpful discussions.

## Conflict of Interest

The authors declare no conflict of interest.

## Author Contributions

R.S.C., P.A.K., A.C., and L.W. conceptualized and designed the experiments. R.S.C. and A.C. conducted all SRS experiments. P.A.K. performed all computational analyses, including DFT calculations, LASSO unmixing, and data visualization. B.S. assisted in neuronal culture. E.D.K. and T.A.T. assisted in performing LC-MS based stable isotope tracing. Z.Y. performed the chemical synthesis experiments. The manuscript was written by R.S.C., P.A.K., A.C., and L.W. with input from all co-authors.

## Notes

### Competing Interest Statement

The authors have declared no competing interest.

