## Supplementary Information for "Optical metabolic imaging of the tricarboxylic acid cycle"

[‡] These authors contributed equally

### Supplementary Figures

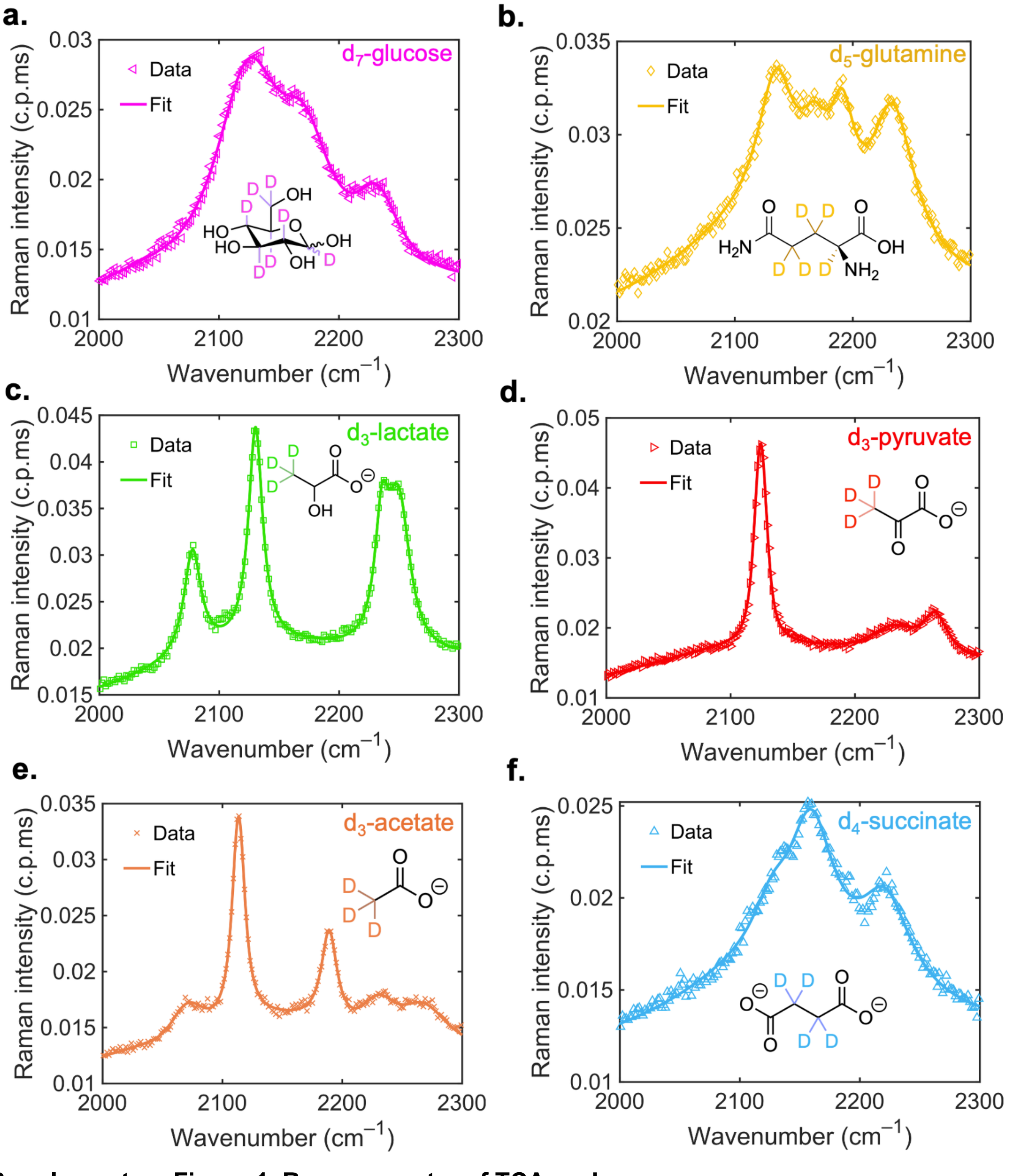

**Supplementary Figure 1. Raman spectra of TCA probes.**

Spontaneous Raman spectra of 200 mM aqueous solutions of **(a)** $d_7$-glucose, **(b)** $d_5$-glutamine, **(c)** $d_3$-lactate, **(d)** $d_3$-pyruvate, **(e)** $d_3$-acetate, and **(f)** $d_4$-succinate (structures inset; c.p.ms, counts per millisecond). A handful of peaks are observed that are not predicted by harmonic DFT calculations (2076 cm$^{-1}$ in $d_3$-lactate; 2070 cm$^{-1}$ and 2190 cm$^{-1}$ in $d_3$-acetate). We attribute these peaks to CD-bending combination bands, similar to previous assignments of Fermi resonances for other deuterated probes.[1]

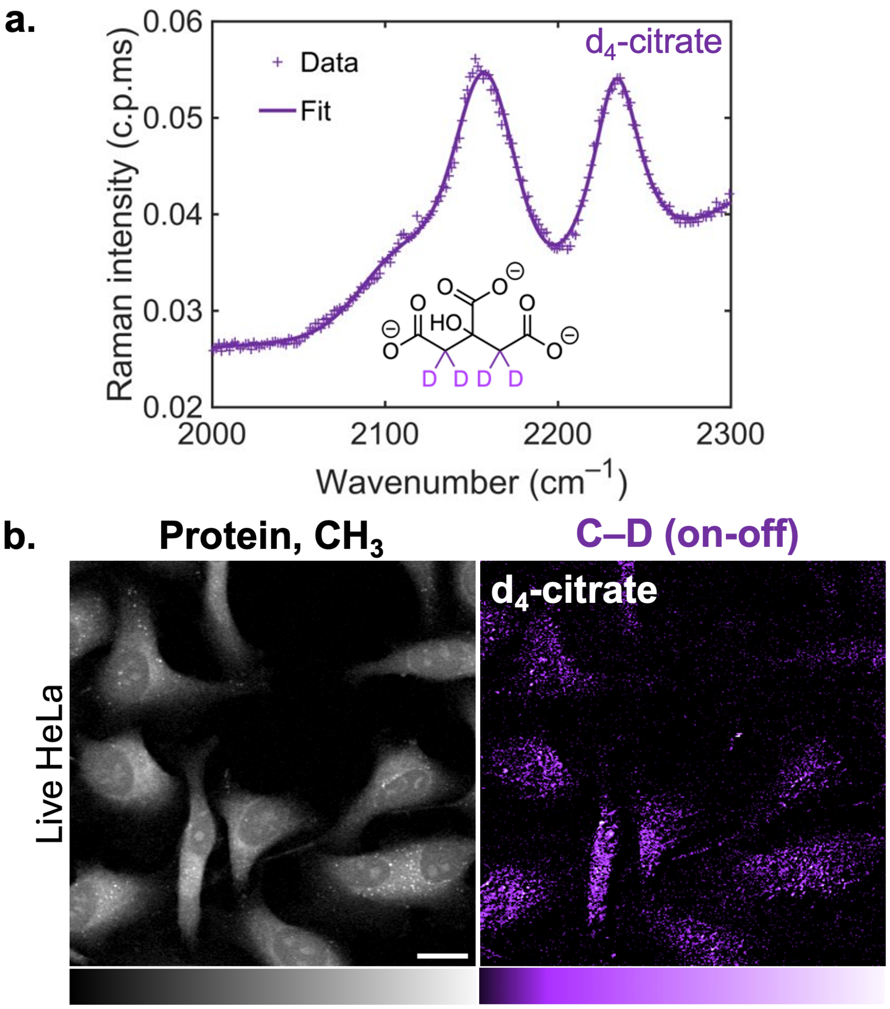

**Supplementary Figure 2. Raman spectra and imaging of $d_4$-citrate.**
**(a)** Spontaneous Raman spectrum of $d_4$-citrate at 1 M in aqueous solution; **(b)** Live-cell SRS imaging of $d_4$-citrate-treated HeLa cells targeted at the protein ($CH_3$, 2940 $cm^{-1}$) and C–D (on-off) channels. Scale bar: 20 μm.

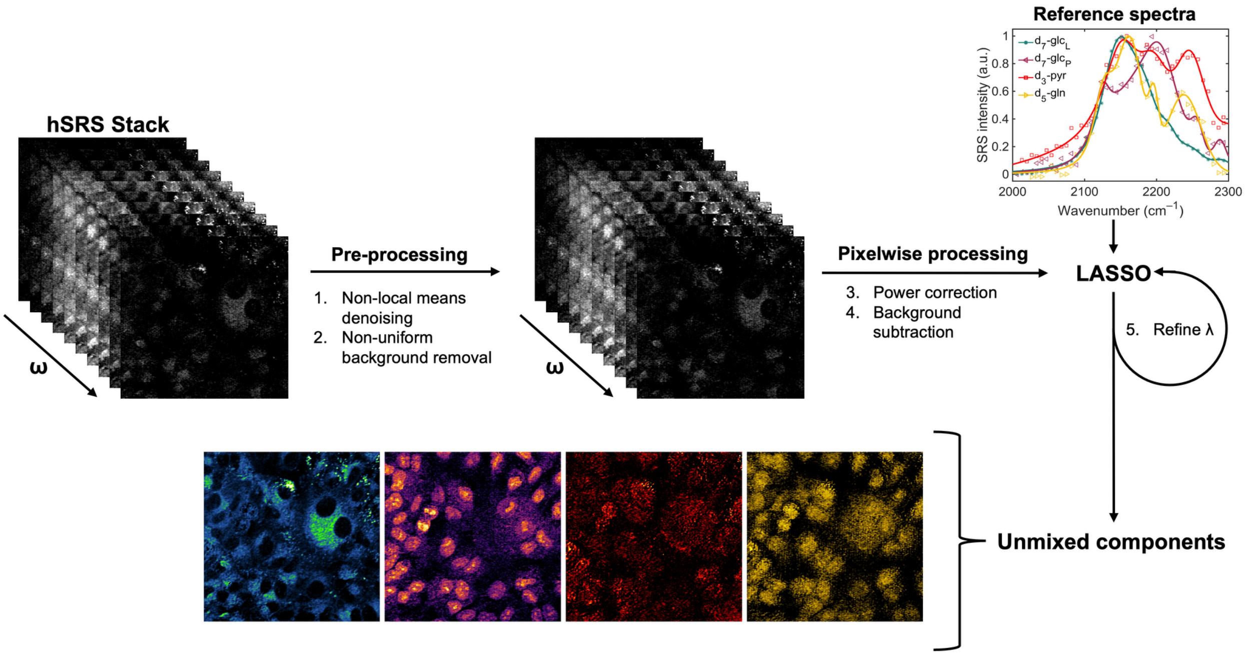

**Supplementary Figure 3. LASSO unmixing workflow.**

Raw hSRS stacks were pre-processed prior to pixelwise power correction, background subtraction, and unmixing via LASSO.

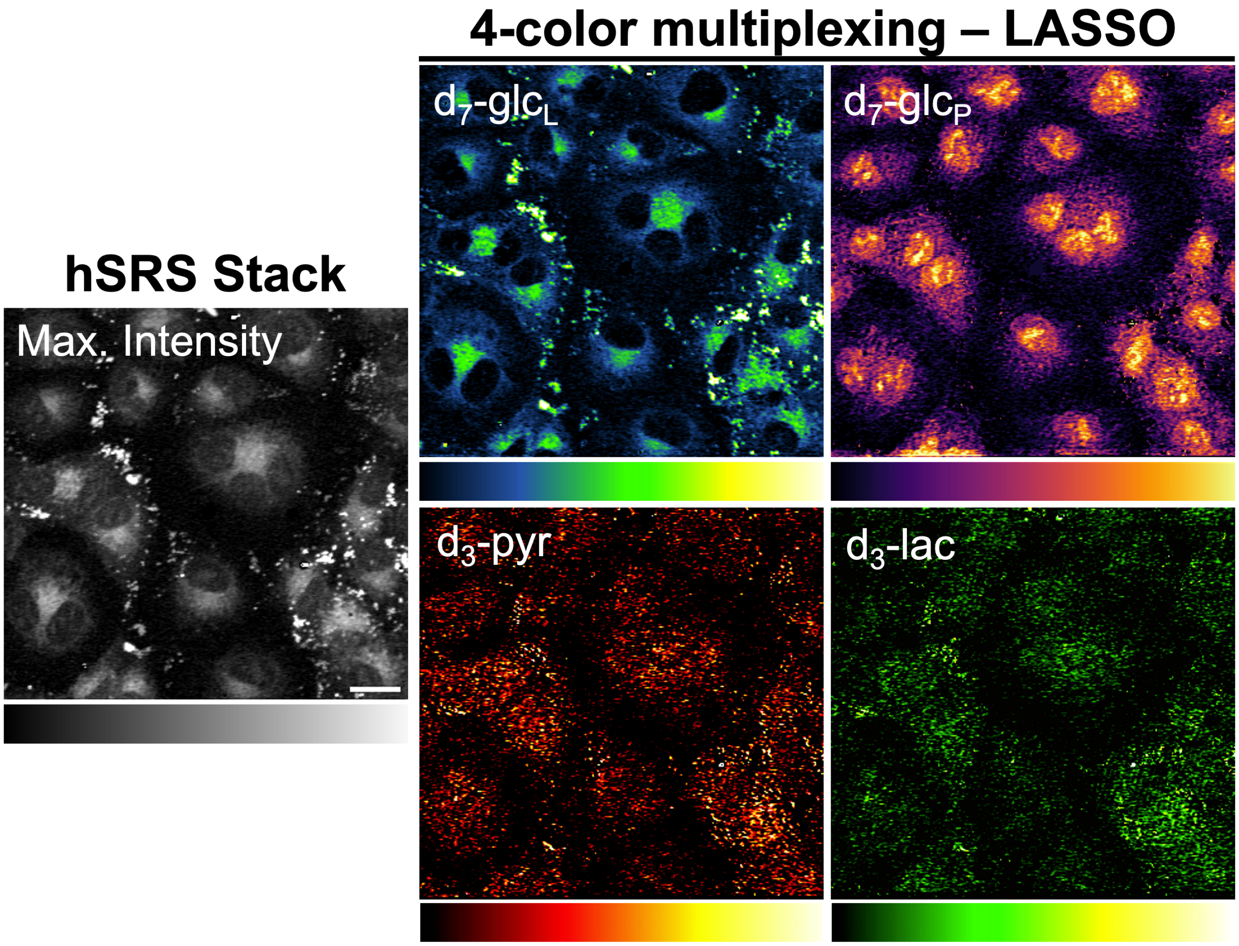

**Supplementary Figure 4. Four-color multiplex imaging with $d_7$-glucose, $d_3$-pyruvate, and $d_3$-lactate.**

Using the same LASSO unmixing pipeline (**Supplementary Figure 3**), we again observe robust unmixing of $d_7$-glc$_L$, $d_7$-glc$_P$, $d_3$-pyr biomass, and $d_3$-lactate biomass. These results serve to demonstrate the broad compatibility of our TCA probes in multiplex imaging with validated chemical unmixing strategies. Scale bar: 20 μm.

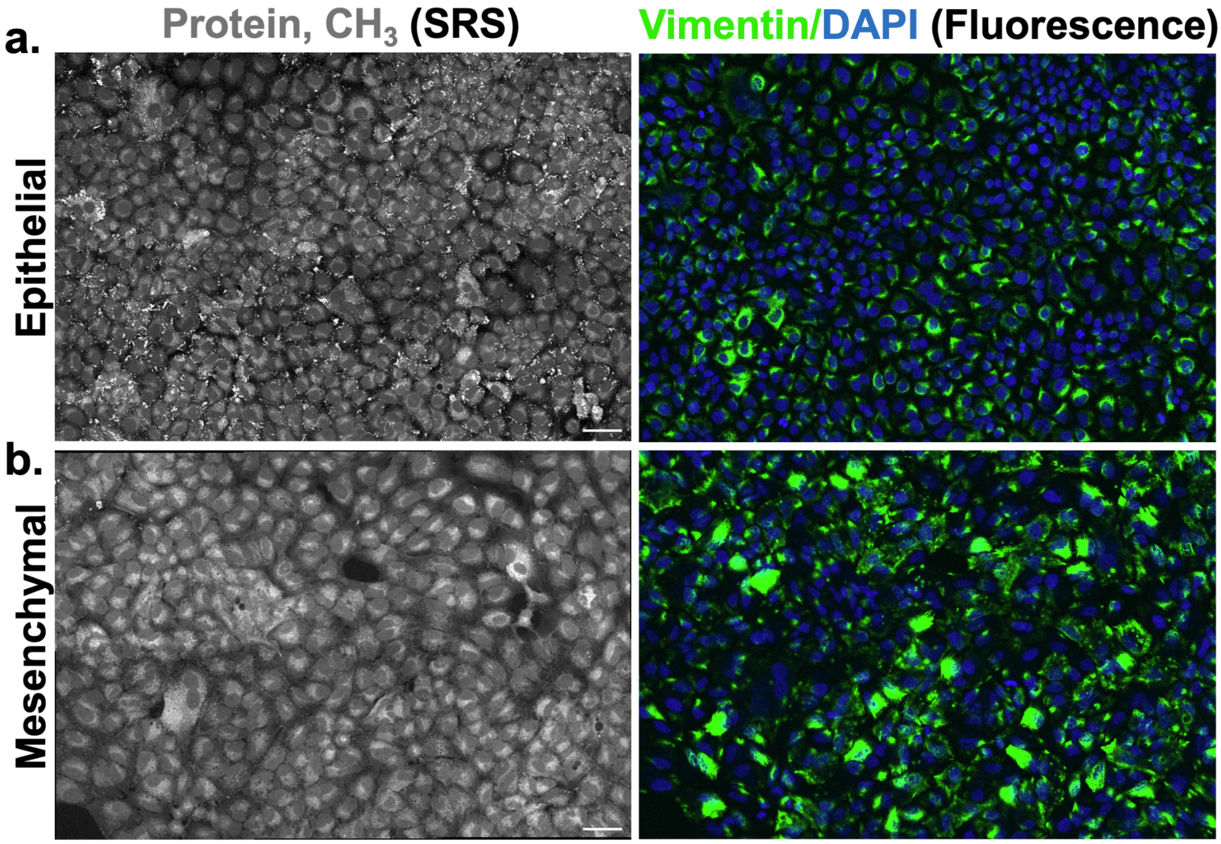

**Supplementary Figure 5. Validation of EMT by immunostaining.**
Correlative SRS (protein channel, 2940 $cm^{-1}$) and Vimentin/DAPI (merged) immunostaining images for **(a)** epithelial and **(b)** mesenchymal cells. Mesenchymal cells exhibit increased vimentin expression. Scale bar: 20 µm.

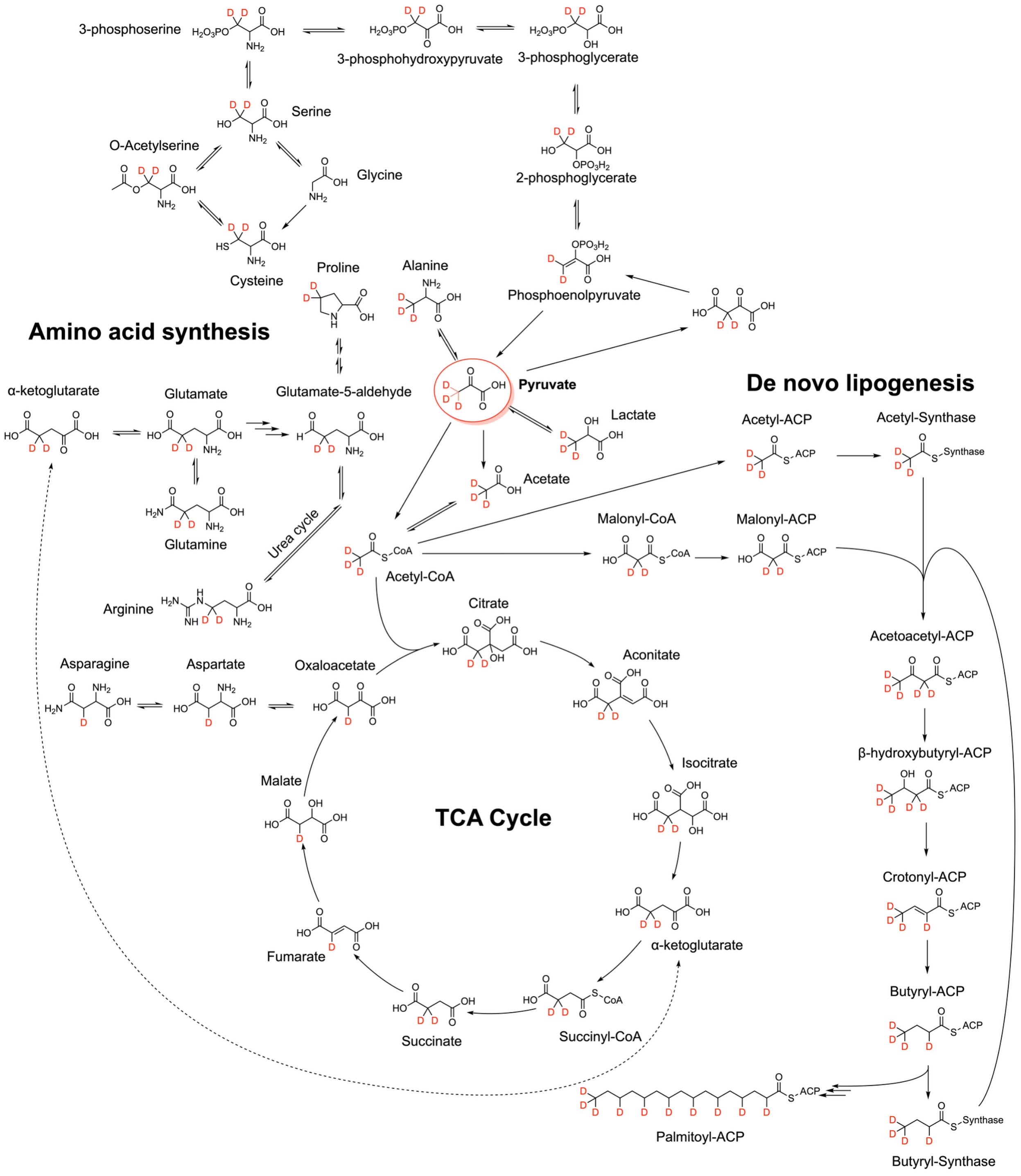

**Supplementary Figure 6. Detailed metabolic reaction mapping.**

$d_3$-pyruvate (red circle) is metabolically incorporated into both amino acids and lipids, with specific structural motifs obtained from the mechanisms of the associated reactions. Here, we show maximal deuteration, and we assume that no deuterium is incorporated from the water in the surrounding media or reducing cofactors, since these would require a massive amount of deuterium in the bulk system to out-compete hydrogen sources.

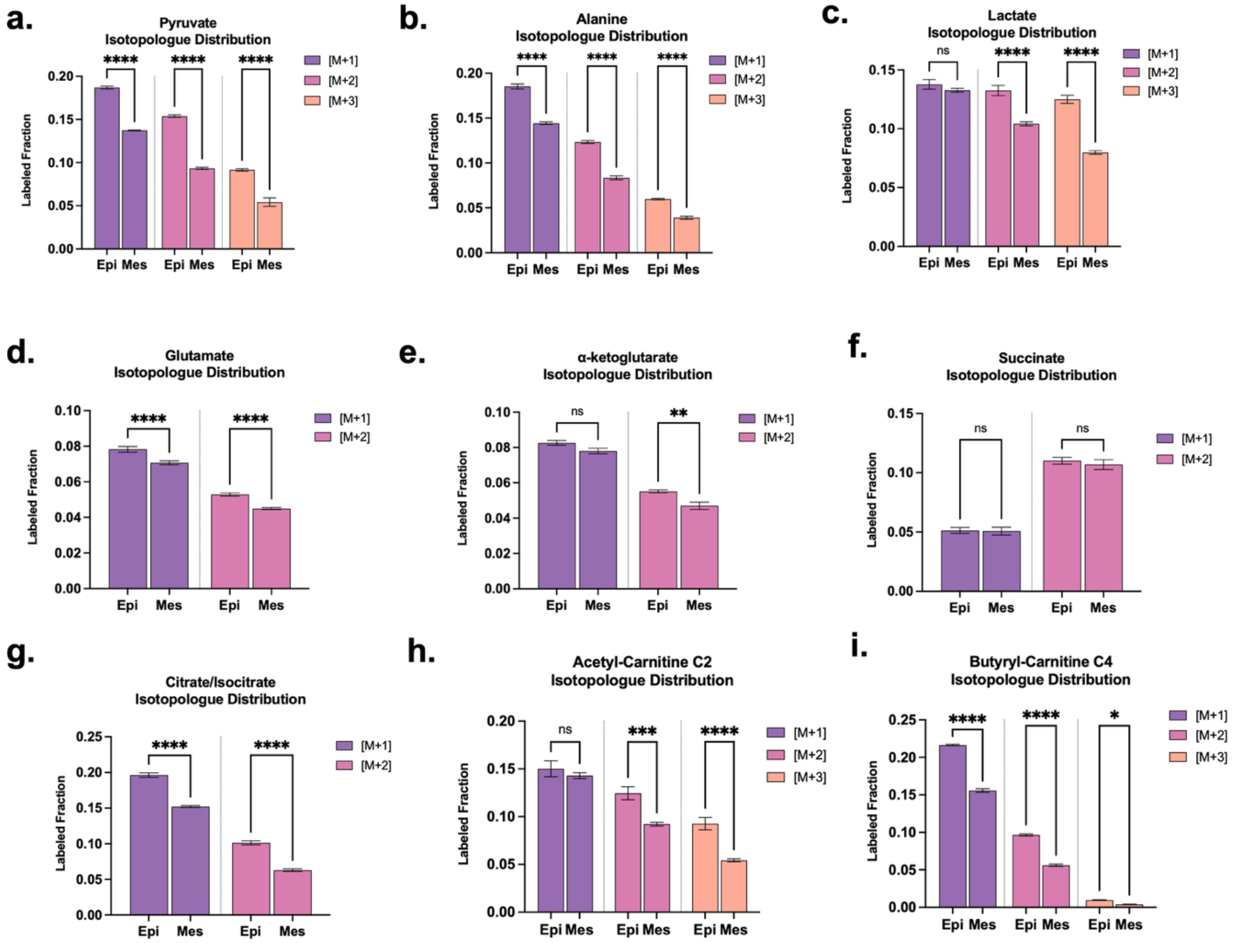

**Supplementary Figure 7. LC-MS quantification of individual metabolites from $d_3$-pyruvate labeling.**

Fractional M+1, M+2, and M+3 isotopologue distributions for **(a)** pyruvate, **(b)** alanine, **(c)** lactate, **(d)** glutamate, **(e)** α-ketoglutarate, **(f)** succinate, **(g)** citrate/isocitrate, **(h)** acetyl-carnitine C2, and **(i)** butyryl-carnitine C4 in epithelial (epi) and mesenchymal (mes) A549 cells. Data from three independent experiments. Data are presented as mean ± SEM from at least three independent experiments. Statistical significance was analyzed using two-way ANOVA test. ****$p < 0.0001$, ***$p < 0.001$, ** $p < 0.01$, *$p < 0.05$, ns = no statistical difference.

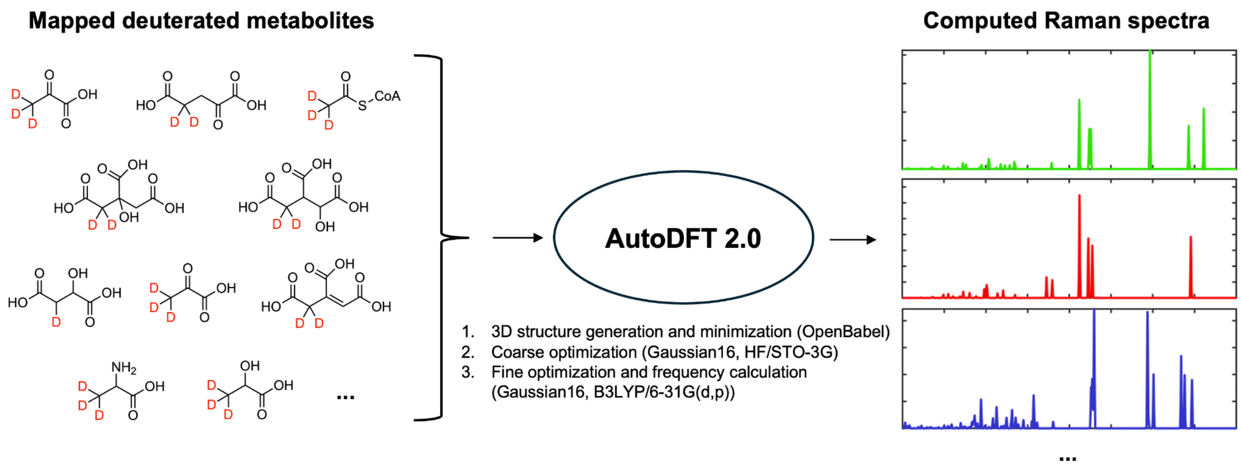

**Supplementary Figure 8. AutoDFT 2.0 workflow.**
Automated density functional theory calculations were carried out with our AutoDFT computational pipeline, with AutoDFT 2.0 now introducing support for selective deuteration and computing Raman scattering activities.

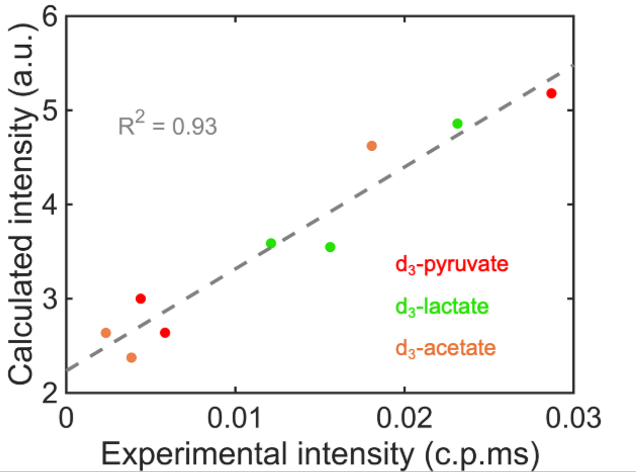

**Supplementary Figure 9. Validation of DFT intensities.**
DFT-calculated Raman spectra were validated by comparison to experimental intensities, obtained by analyzing the spectra shown in **Figure S1**. Only the $d_3$ species were used for intensity validation because they exhibited distinct, well-separated bands that allowed confident quantitation. For the more highly deuterated probes, highly overlapping and broad peaks prohibited confident experimental peak quantification (fits were sensitive to initial conditions and yielded inconsistent intensities, while the frequencies were more robust).

We expect this to have a minimal impact on the quantitative analysis presented in the main text (e.g., **Fig. 5d**), since the internal consistency of DFT means that this accuracy should generalize to more than simple $d_3$ species. Additionally, since we apply our spectral profiling only to four well-resolved spectral bands (2120, 2160, 2200, and 2240 $cm^{-1}$), our quantitation of deuterated biomass is expected to be more robust.

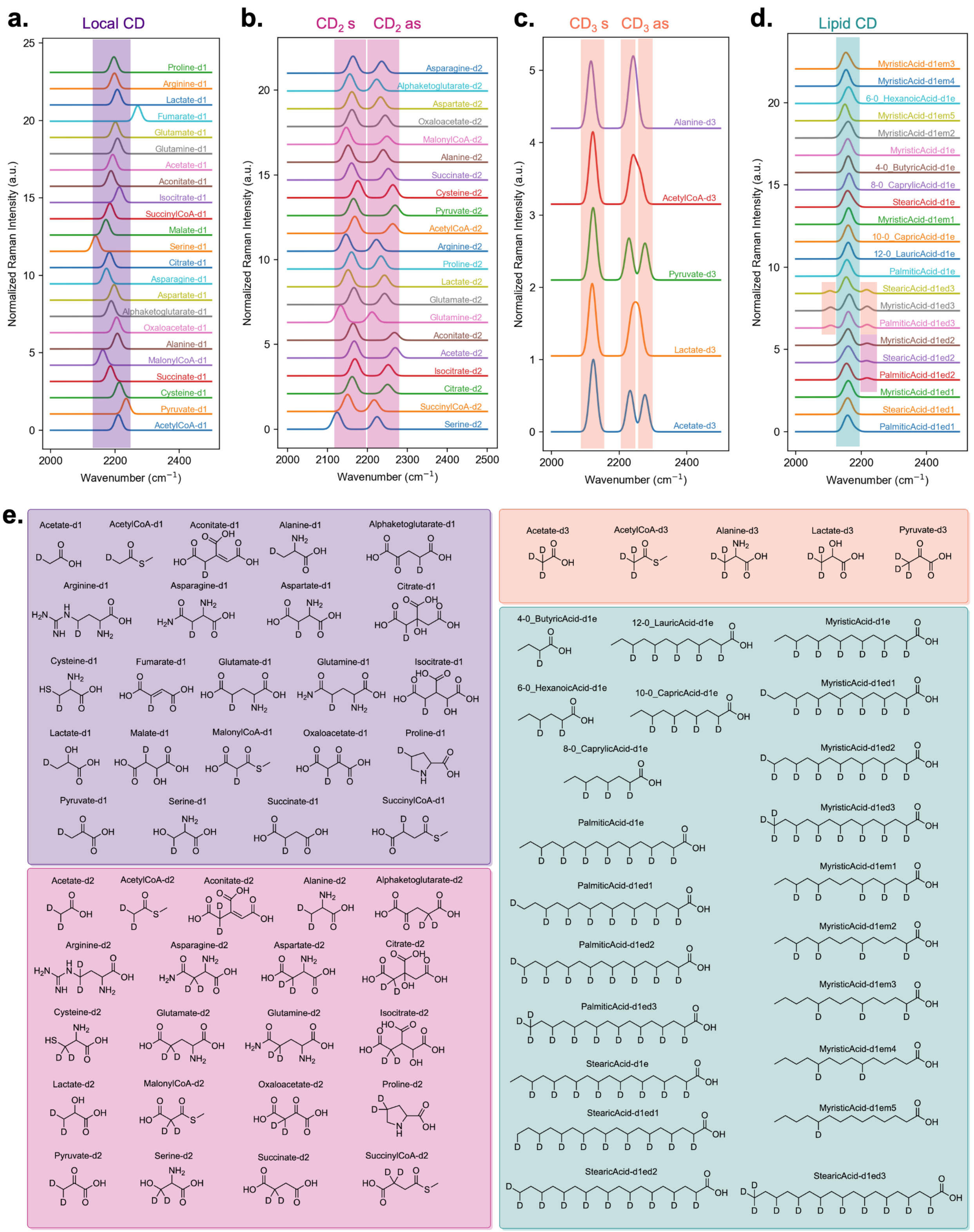

**Supplementary Figure 10. DFT-computed spectra of mapped metabolites from $d_3$-pyr.**
**(a)** Computed spectra of $d_1$ metabolites, showing a local C–D mode at ~2200 $cm^{-1}$ (purple shaded area). The main exception is $d_1$-fumarate, which contains a $C_{sp2}$–D bond and is blueshifted relative

to the $C_{sp3}$–D local stretching band at 2200 $cm^{-1}$; **(b)** Computed spectra of $d_2$ metabolites, showing symmetric ($CD_2$ s) and asymmetric ($CD_2$ as) stretching bands at ~2160 and ~2240 $cm^{-1}$, respectively (pink shaded areas); **(c)** Computed spectra of $d_3$ metabolites, showing a symmetric stretching band ($CD_3$ s) at ~2120 $cm^{-1}$ and two asymmetric stretching bands ($CD_3$ as) at ~2230 and ~2250 $cm^{-1}$ (orange shaded areas), often clustering into a single band at ~2240 $cm^{-1}$; **(d)** Computed spectra of aliphatic chains ($CDHCH_2$), displaying a coupled CD band at ~2160 $cm^{-1}$ (teal shaded area). For lipids with $d_2$ and $d_3$ methyl groups, the symmetric and asymmetric bands of the methyl groups are observed at the same characteristic frequencies as the $d_2$ and $d_3$ metabolites (highlighted regions in orange and pink); **(e)** Chemical structures of all calculated metabolites, grouped by deuteration ($d_1$, purple; $d_2$, pink; $d_3$, orange; fatty acids, teal). CoA units were truncated and replaced by methyl substituents. All DFT calculations were performed on neutral molecules (as drawn). Mapped metabolites are named by their deuteration status, broadly grouped as $d_1$, $d_2$, $d_3$, and aliphatic chains. The aliphatic chains all exhibit the $(CDHCH_2)_n$ deuteration motif, which we derived from our reaction network mapping (**Supplementary Figure 6**) and label here as “d1e” in shorthand, indicating a single deuterium on every other carbon in the chain. To account for the large potential heterogeneity of lipid biomass, we varied the chain length (ranging from 4 carbons to 18 in increments of 2) and the deuteration of the terminal methyl group (labeled “d1e**d1**,” d1e**d2**,” or “d1e**d3**”). We also explored the effects of missing deuteriums across a set of myristic acid structures (labeled “d1em#,” where # is the number of deuteriums missing, relative to the “d1e” structure).

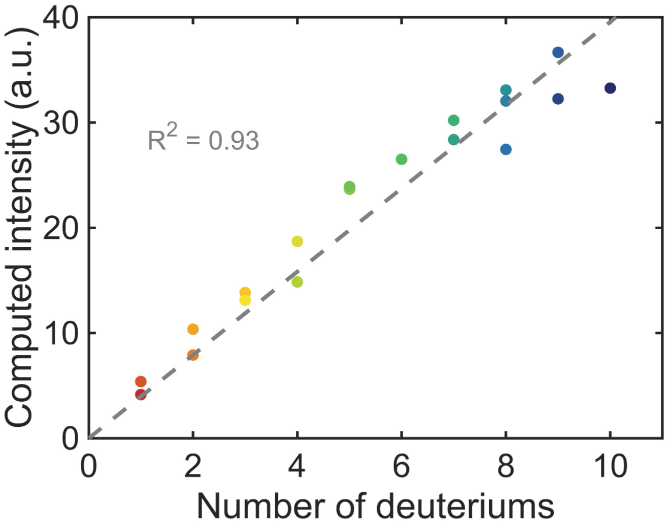

**Supplementary Figure 11. Lipid computed cross-sections as a function of deuterium content.**

The calculated Raman scattering cross-sections for mapped lipids ($CDHCH_2$) are strongly linearly correlated with the number of deuteriums in the lipid, allowing us to quantify total deuterium in aliphatic chains (at 2160 cm$^{-1}$) on a per-deuterium basis.

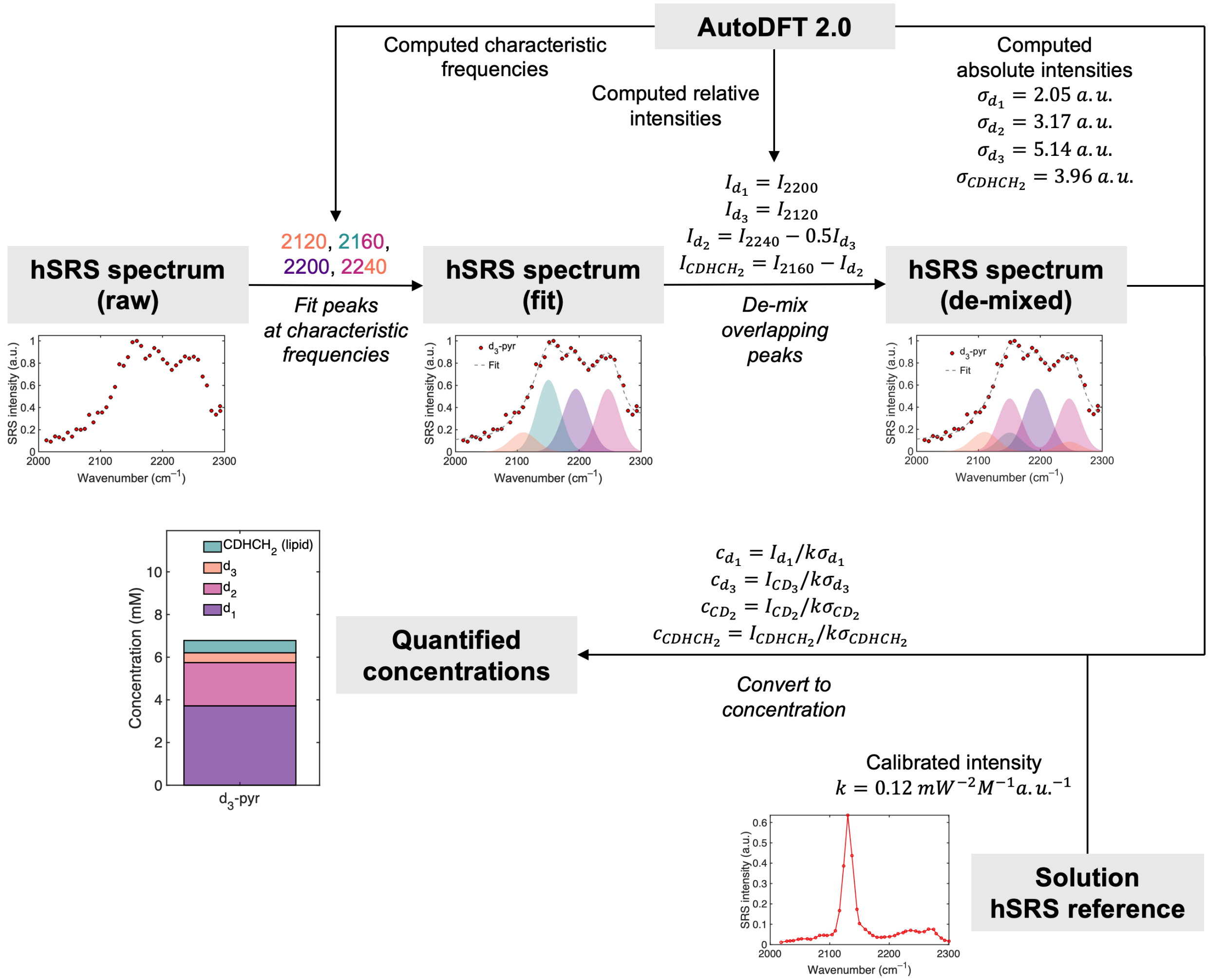

**Supplementary Figure 12. DFT-derived spectral analysis workflow.**
The obtained hSRS spectrum from a given sample is fit to four Gaussian peaks at the characteristic frequencies obtained by DFT (allowed to float within ± 10-15 cm$^{-1}$). The fit amplitudes ($I_{2200}$, etc.) at each frequency are de-mixed into amplitudes for each metabolite class ($I_{d_1}$, etc.) by accounting for spectral mixing (assuming a 1:2 ratio for $d_3$ species at 2120 and 2240 cm$^{-1}$ and a 1:1 ratio for $d_2$ species at 2160 and 2240 cm$^{-1}$). De-mixed amplitudes are then converted to abundances by dividing by the mean cross-sections of each metabolite class ($\sigma$) and converted to absolute concentrations from a solution reference measurement of $d_3$-pyruvate in water ($k$). To estimate error, we calculated the root-mean-square error in intensity in our DFT validation (**Supplementary Figure 9**), which is 0.29 a.u. (corresponding to 6-15% error). Taking this error as σ and propagating through the conversion to concentration yields an estimated 30% (2σ) error in SRS-derived quantification, which we use for all derived concentrations.

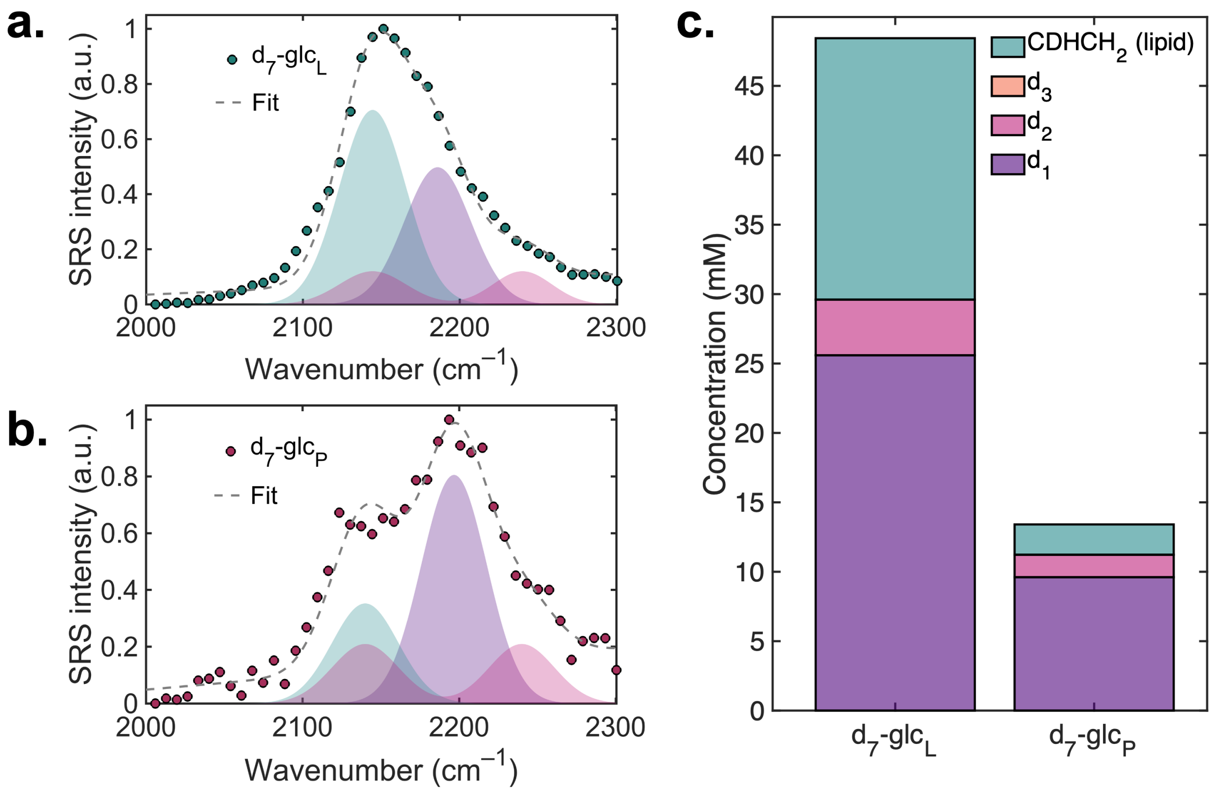

**Supplementary Figure 13. SRS quantification of $d_7$-glucose-derived biomass.**
Fitting of hSRS **(a)** $d_7$-glc$_L$ and **(b)** $d_7$-glc$_P$ spectra. **(c)** Quantification of $d_1$, $d_2$, and $CDHCH_2$ biomass. Glucose-derived biomass exhibits impressively high total concentrations of 48±15 mM in $d_7$-glc$_L$ and 13±4 mM in $d_7$-glc$_P$ (spectra obtained from lipid droplets and nucleoli, respectively; concentrations tabulated in **Supplementary Table 1**), despite the cell media containing only 25 mM $d_7$-glc. However, considering that $d_7$-glc contains multiple deuterium equivalents per molecule (i.e., through glycolysis, two molecules of $d_2$-pyruvate are generated per molecule of $d_7$-glc, setting an estimated upper-bound at 50 mM), these results appear quite reasonable, with near-maximal incorporation efficiency for $d_7$-glc in lipid droplets.

With our DFT-derived spectral profiling, we can additionally examine the distributions of $d_1$, $d_2$, $d_3$ and lipid ($CDHCH_2$) deuterium in each of the spectra. Both $d_7$-glc$_L$ and $d_7$-glc$_P$ exhibit no $d_3$ concentration, since $d_7$-glc contains no deuterated methyl groups. The $d_7$-glc$_P$ spectrum is dominated by $d_1$ species (10±3 mM), indicating that nucleolar proteins are synthesized relatively slowly compared with deuterium abstraction (i.e., $d_3 \rightarrow d_2 \rightarrow d_1$), as by the TCA cycle and other related reactions (**Supplementary Figure 6**). More interestingly, our lipid droplet-derived biomass ($d_7$-glc$_L$) exhibits a $d_1$ concentration of 26±8 mM (which is spectrally distinct from lipid-associated deuterium, even at the singly deuterated level; **Supplementary Figure 10**). Although some of this signal results from single deuteration of the terminal methyl group, the $d_1$ concentration exceeds that of aliphatic deuterium in lipid droplets (19±6 mM). These results suggest that a significant amount of deuterium biomass in lipid droplets may in fact originate from phospholipid head groups (e.g., serine or choline) or lipid droplet-associated proteins, such as the perilipin family, rather than comprising purely fatty acid-associated deuterium. Further work is needed to fully understand these results.

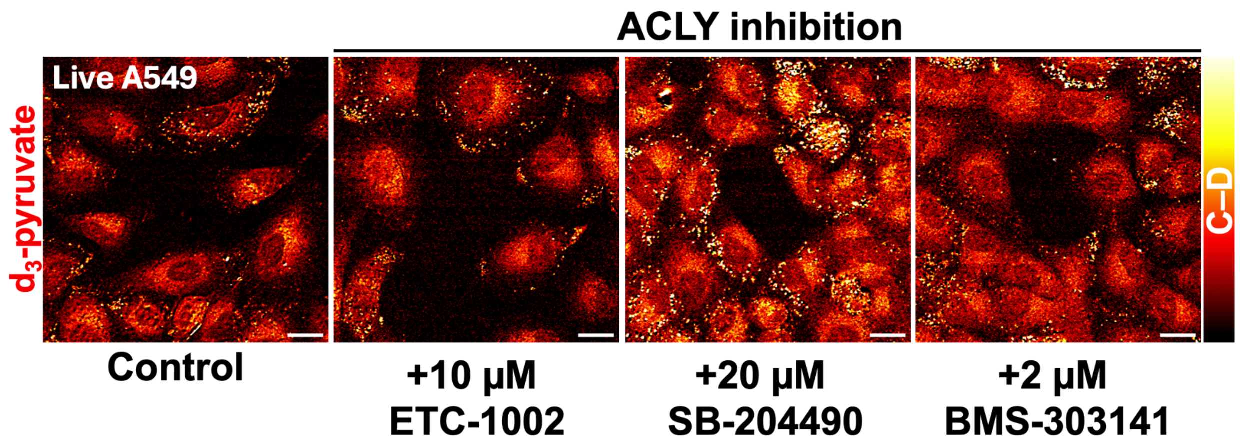

**Supplementary Figure 14. ACLY inhibition of $d_3$-pyruvate metabolism in live A549 cells.** Inhibition of ACLY via three separate drugs (ETC-1002, SB-204490, and BMS-303141) yielded no obvious attenuation of $d_3$-pyruvate biomass in A549 cells, which is consistent with $d_3$-pyruvate shuttling into lipids via ACSS2 (bypassing the inhibited pathway). Scale bar: 20 μm.

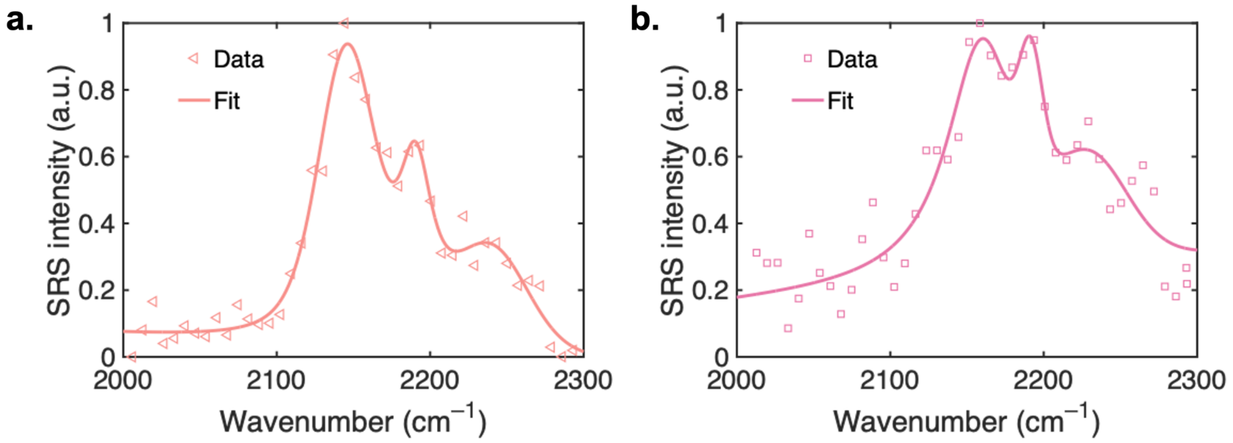

**Supplementary Figure 15. Pyruvate biomass spectra under pharmacological perturbation.** **(a)** hSRS spectrum of the rigid solid membrane (SM) obtained with inhibition of SCD1 with CAY10566; **(b)** hSRS spectrum of total cytoplasmic biomass obtained with upregulation of protein by CHIR99021.

### Supplementary Tables

**Supplementary Table 1. Quantified concentrations from biomass spectra.**

| Conc. (mM) | $d_1$ | $d_2$ | $d_3$ | $CDHCH_2$ | Total |
|---|---|---|---|---|---|
| **$d_7$-glc$_L$** | 25.6 ± 7.7 | 4.0 ± 1.2 | 0 | 18.8 ± 5.6 | 48.4 ± 14.5 |
| **$d_7$-glc$_P$** | 9.6 ± 2.9 | 1.6 ± 0.5 | 0 | 2.2 ± 0.7 | 13.4 ± 4.0 |
| **$d_3$-pyr** | 3.7 ± 1.1 | 2.0 ± 0.7 | 0.5 ± 0.1 | 0.6 ± 0.2 | 6.8 ± 2.0 |
| **$d_3$-pyr SM** | 9.2 ± 2.8 | 2.4 ± 0.7 | 1.2 ± 0.4 | 6.0 ± 1.8 | 18.9 ± 5.7 |
| **$d_3$-pyr+CHIR** | 7.0 ± 2.1 | 1.0 ± 0.3 | 1.6 ± 0.5 | 2.1 ± 0.6 | 11.7 ± 3.5 |

### Supplementary Materials & Methods

#### Preparation and imaging of $d_4$-citrate.

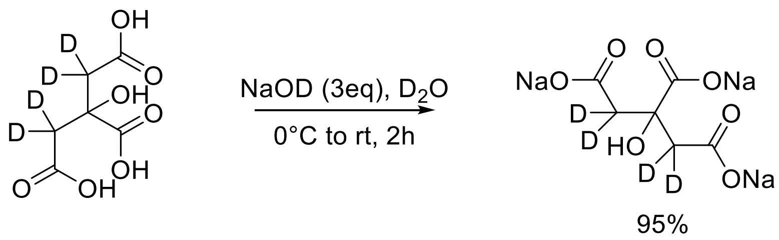

For metabolic imaging of $d_4$-citrate, we first formed a trisodium salt using commercially available 2,2,4,4-$d_4$-citric acid (1.92 g, 10.0 mmol; Cambridge Isotope, Cat. #DLM-3487-0.5) that was added to a flame-dried flask in 10 mL deuterium oxide as the solvent. Following this, NaOD 50% mass/$D_2O$ solution (2.4 g, 30.0 mmol) was slowly added to the reaction mixture on an ice bath. The reaction was then allowed to react at room temperature for 2h and its completion was monitored using H-NMR. Upon full conversion, the solvent was evaporated with a rotatory evaporator to obtain the final product $d_4$-citrate-trisodium salt, as a white solid with 95% yield.

For metabolic labeling, HeLa cells were incubated with 30 mM $d_4$-citrate for 48 hours before SRS imaging.
